# RecV recombinase system for *in vivo* targeted optogenomic modifications of single cells or cell populations

**DOI:** 10.1101/553271

**Authors:** Shenqin Yao, Peng Yuan, Ben Ouellette, Thomas Zhou, Marty Mortrud, Pooja Balaram, Soumya Chatterjee, Yun Wang, Tanya L. Daigle, Bosiljka Tasic, Xiuli Kuang, Hui Gong, Qingming Luo, Shaoqun Zeng, Andrew Curtright, Ajay Dhaka, Anat Kahan, Viviana Gradinaru, Radosław Chrapkiewicz, Mark Schnitzer, Hongkui Zeng, Ali Cetin

## Abstract

Brain circuits are composed of vast numbers of intricately interconnected neurons with diverse molecular, anatomical and physiological properties. To allow highly specific “user-defined” targeting of individual neurons for structural and functional studies, we modified three site-specific DNA recombinases, Cre, Dre and Flp, by combining them with a fungal light-inducible protein, Vivid, to create light-inducible recombinases (named RecV). We generated viral vectors to express these light-inducible recombinases and demonstrated that they can induce genomic modifications in dense or sparse populations of neurons in superficial as well as deep brain areas of live mouse brains by one-photon or two-photon light induction. These light-inducible recombinases can produce highly targeted, sparse and strong labeling of individual neurons in multiple loci and species. They can be used in combination with other genetic strategies to achieve specific intersectional targeting of mouse cortical layer 5 or inhibitory somatostatin neurons. In mouse cortex sparse light-induced recombination allows whole-brain morphological reconstructions to identify axonal projection specificity. Furthermore these enzymes allow single cell targeted genetic modifications via soma restricted two-photon light stimulation in individual cortical neurons and can be used in combination with functional optical indicators with minimal interference. In summary, RecVs enable spatiotemporally-precise, targeted optogenomic modifications that could greatly facilitate detailed analysis of neural circuits at the single cell level by linking genetic identity, morphology, connectivity and function.

## INTRODUCTION

To understand how a biological system works it is often useful to define its building blocks and understand how those building blocks work together to generate its function. Mammalian brain is one of the most complex biological systems. It is composed of millions to billions of cells^1^ with diverse characteristics. To understand this extraordinary complexity, it will be essential to define cell types based on properties such as gene expression, morphology and physiology at the single cell level. Furthermore, the unique properties of individual cells need to be related to their connectivity patterns and their activities in a behavioral context. Anatomical information combined with genetic identity and functional properties at the single cell level will enable better analysis of brain circuitry underlying complex behaviors in health and disease.

One of the most powerful approaches to characterizing cell types and studying their functions relies on mouse genetics^2^. Strategies that rely on transgenic or viral expression of recombinases allow a highly specific level of genetic modification^3–5^. A multitude of cell type-specific Cre recombinase mouse lines has led to many discoveries in neuroscience over the last few decades. Further improvements on spatiotemporal control can allow much higher resolution manipulations and studies of biological systems.

Currently finding individual cells/neurons *in vivo*; characterizing and genetically manipulating them for targeted genomic modifications as well as expressing genes in a targeted manner is challenging to execute in an efficient manner. The state-of-the-art approach for introducing an exogenous gene to a specific neuron is either by the patch clamp technique^6^ or via single-cell electroporation^7,8^. These techniques are highly challenging and usually result in low and variable yields. Sparse neuronal labeling or manipulation can be achieved by controlling transgene recombination by lowering the dose of inducers (*e.g.*, tamoxifen, in the case of CreER) or by employing ‘inefficient’ recombinase reporters (*e.g.*, MADM^9^). However, the sparse genetic modification achieved using these methods is random and cannot be easily directed to specific or individual cells of interest.

Light is a particularly powerful and versatile regulator with its tremendous adjustability in the dimensions of spectrum, intensity, space (location and size) and time (timing and duration). Using photons to access and genetically modify individual neurons will offer an improvement over the current state-of-the-art. Multi-photon additive characteristics of light that can generate a spatiotemporally-restricted excitation are well suited for achieving fine spatial control^10^. Thus, modifying current genomic manipulation enzymes to make them light inducible could be an ideal approach to reach a high spatiotemporal resolution for targeted single cell manipulations.

Several light inducible protein-based spatiotemporal control systems have been developed to date^11–33^. These systems depend on various light sensing modules to control protein states, protein localization, transcription and genetic alterations. So far optical manipulation of genomes -optogenomics-of individually targeted single cells within live intact tissues has not been demonstrated using such methods. To leave a permanent genetic mark in individual cells in this study, we developed and validated light inducible site-specific recombinase systems and its associated set of viral tools via comparing and optimizing various light inducible systems with the criteria that they induce robust genomic modifications with no or minimally detectable background under no-light conditions at different genomic locations and species. Our work resulted in the generation of highly efficient light inducible versions of the most commonly used site-specific recombinases-Cre, Dre and Flp - that allow tight population level or target-specific single cell level optogenomic modifications *in vivo*.

## RESULTS

### A Split Vivid-Cre enables efficient light-inducible site-specific DNA modifications

To spatiotemporally regulate site-specific recombination, we generated a light-inducible genetic switch based on a fungal light sensitive protein – Vivid (VVD)^34^. We chose VVD mainly because it is the smallest (450 bp) of all Light Oxygen or Voltage (LOV) domain-containing proteins, and therefore suitable for fitting into viral vectors with limited genomic capacity. In addition, the spectral properties of VVD -its excitation and emission drops sharply to near 0 with light above ~520 nm- would allow us to use other fluorescent proteins, activity indicators or optogenetic molecules that work at longer wavelengths^20,35^.

Upon light illumination, VVD forms a homodimer^35^ due to the conformational changes induced by Flavin Adenine Dinucleotide (FAD) cofactor within the LOV domain. The FAD cofactor within the LOV domain has a peak single-photon (1P) excitation at 450 nm and a peak two-photon (2P) activation at 900 nm wavelength^36^.

To generate a precise spatiotemporal control of gene expression, we used a split-Cre recombinase system^20,37–39^ and converted it into a light-inducible system using the VVD protein (**Fig. 1a**). For Cre recombinase to function properly, the N-terminal and the C-terminal portions of the protein need to come in close proximity to form the correct tertiary structure. The crystal structure of Cre is known^40^ and so of VVD^35,41^. In the dimer form, the N-terminus of one VVD monomer gets into close proximity to the C-terminus of the other (**Fig. 1a**). Guided by this information our design attempted to bring together the N- and C-portions of Cre in the correct orientation upon light-induced conformational change and dimerization of VVD. To split the Cre protein, we relied on a previous split Cre system, since it was shown to generate stable dimers^37^. We fused the inactive N-terminal segment of Cre to the N terminus of one VVD monomer and inactive C- terminal segment of Cre to the C terminus of another VVD monomer, which we codon diversified to reduce the sequence similarity between the two VVD monomers and minimize the risk of recombination. We cloned each of these into recombinant adeno-associated virus (rAAV) expression vectors (**Fig. 1c**) ^42^. The resulting NCreV and CCreV constructs along with fluorescent Cre reporter constructs were co-transfected into mammalian cells and robust light-inducible recombination was observed with 458 nm LED light induction compared to the no-light conditions (**Fig. 1d, e**). We named these new proteins CreV, and the general light-inducible Vivid-recombinase system RecV.

**Figure 1.**
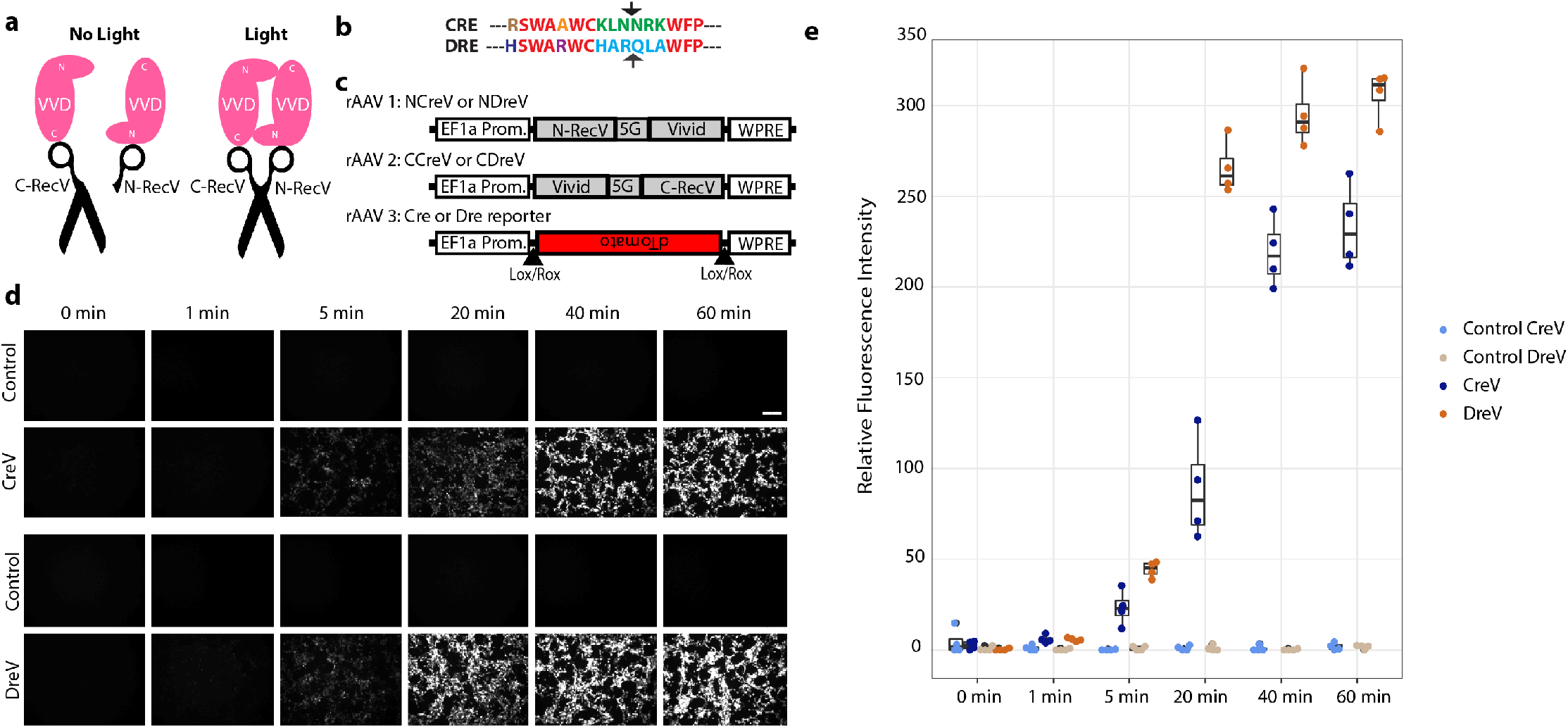
Design of the RecV systems. (**a**) The N terminus of one VVD protein gets in close proximity to the C terminus of the second VVD protein in dimer form based on the crystal structure. Cre is split into two portions: N-Cre and C-Cre. Upon light induced dimerization of the VVD proteins, the two portions of Cre are brought together and Cre activity is restored (right panel). (**b**) Alignment of amino-acid sequences of Cre and Dre recombinases, focusing on the location for the split (arrows). Using this information, we split Dre at the arginine to glutamine (N) transition marked with the arrow. (**c**) Schematic of the CreV and DreV rAAV constructs. CreV (NCreV + CCreV) or DreV (NDreV + CDreV) fusion constructs were co-transfected with a Cre- or Dre- dependent red fluorescent reporter. (**d**) Results of different durations of light induction at 458 nm wavelength of a 1.3 mW/mm^2^ LED light source were documented at 48 hours post induction, or dark as controls. Scale bar (top right), 200 μm. (**e**) Quantification of relative fluorescence intensity of the reporter constructs shows light-induced recombination for by CreV or DreV depending on the duration of light stimulation. Each experiment is represented by 4 replicates.

### Vivid-based light-inducible system for Dre recombinase

To broaden the capabilities of the RecV approach we modified another site-specific DNA recombinase, Dre ^43^, which is a relative of Cre, and is likewise derived from a bacterial phage. Dre recombinase recognizes a sequence called Rox^44^, which is different from LoxP. This is especially important since it would allow us to utilize already existing Cre driver mouse lines to study Cre-defined populations and to develop intersectional strategies to further refine cell type-specific genetic manipulation.

We reasoned that the sequences surrounding the homology region within Dre N-terminus, where we split Cre into two non-functional compartments, could also be targeted to generate a split Dre (**Fig. 1b**). After generation of the DreV constructs based on CreV designs we tested the light inducibility of DreV constructs using fluorescent Dre reporter plasmids (**Fig. 1c** and **Supp. Fig. 1**). Our results indicate that the light-induced DreV recombination is efficient using 1P excitation and is similar to CreV (**Fig. 1d, e**).

Spatiotemporal regulation of gene expression can be further fine-tuned using 2P excitation since it has a much narrower point spread function in the axial direction. We tested if the VVD based recombination can take place using 2P light under cell culture conditions. Our results indicate that DreV-mediated light-inducible recombination can be achieved by application of 900-nm wavelength of 2P light (**Supp. Fig. 2**).

### Single RecV expression constructs and comparison to other existing light inducible recombinase systems

To implement the RecV strategy efficiently and reduce the number of viruses or transgenic mouse lines needed to use this system, it is preferable to co-express the two halves of RecVs in a single construct. To achieve this, we tested a variety of ways to link the N and C components of CreV or DreV that would provide the highest efficiency of recombination with the least amount of background recombination: IRES, 2A, single open reading frame with permutations of the open reading frames; see methods section for more details (**Supp. Fig. 3a**). Among these constructs highest efficiencies were observed when optimized elements from Cre-Magnets (mutant VVDs)^13,33^ were used, which we named iCreV and iDreV. iCreV and iDreV are composed of N- and C-terminal portions of the Cre and Dre split recombinases fused to nuclear localized wild-type VVD and co-expressed with optimized linkers and 2A elements that were used in Cre-Magnets (**Supp. Fig. 1**).

**Figure 2.**
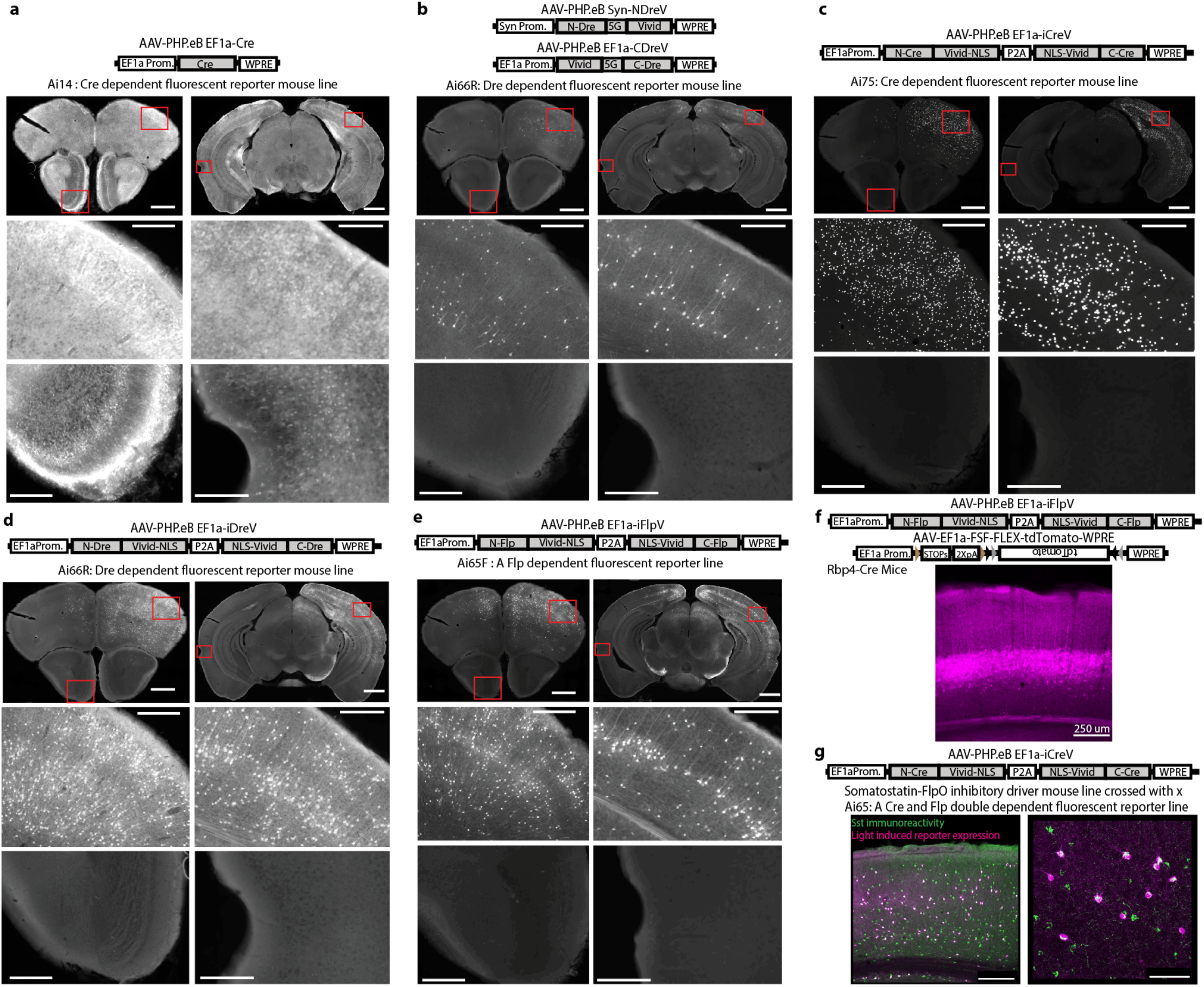
Optogenomic modifications by RecV viruses with spatiotemporal and intersectional cell class specific precision *in vivo*. Reporter mice (n = 2 per case) received right hemisphere intracerebroventricular (ICV) or retroorbital (RO) injection of PHP.eB rAAVs followed by left hemisphere light stimulation 2-weeks post injection. Brains were examined by fluorescence microscopy 2 weeks post-light stimulation. (**a**) Cre-dependent tdTomato-expressing Ai14 mice were ICV injected with AAV-PHP.eB EF1a-Cre virus leading to widespread recombination throughout the brain. (**b**) Dre-dependent tdTomato-expressing Ai66R mice were ICV injected with a 1:1 mixture of AAV-PHP.eB Syn-NDreV and AAV-PHP.eB EF1a-CDreV. (**c**) Cre-dependent nls-tdTomato expressing Ai75 mice were ICV injected with AAV-PHP.eB EF1a-iCreV. (**d**) Ai66R mice were ICV injected with AAV-PHP.eB EF1a-iDreV. (**e**) Flp-dependent tdTomato-expressing Ai65F mice were injected with AAV-PHP.eB EF1a iFlpV. Scale bars a-e, 1 mm for top images, 200 μm for bottom images. (**f**) Layer 5 pyramidal neuron-specific Rbp4-Cre mice were injected with a mixture of AAV-PHP.eB EF1a-iFlpV and AAV-PHP.eB EF1a-FSF-FLEX-tdTomato. Scale bar, 250 μm. (**g**) Somatostatin FlpO mouse line, Sst-IRES-FlpO, crossed with a Cre/Flp double dependent fluorescent reporter mouse line, Ai65, was RO injected with AAV-PHP.eB EF1a-iCreV and light was delivered on the left hemisphere. Specific intersectional light induced recombination was observed in somatostatin positive inhibitory interneurons as revealed by immunohistochemistry. Scale bar, 250 μm for left and 75 μm for right images. All *in vivo* light activation was applied through the skull on the left hemisphere (opposite of the ICV injection sites in those cases). For a-e, two coronal planes are shown for each injection (top row) with enlarged views (lower two rows) for areas indicated by the red boxes. In all experimental cases (**b-g**) light induction led to localized recombination radiating from the light induced hemisphere.

**Figure 3.**
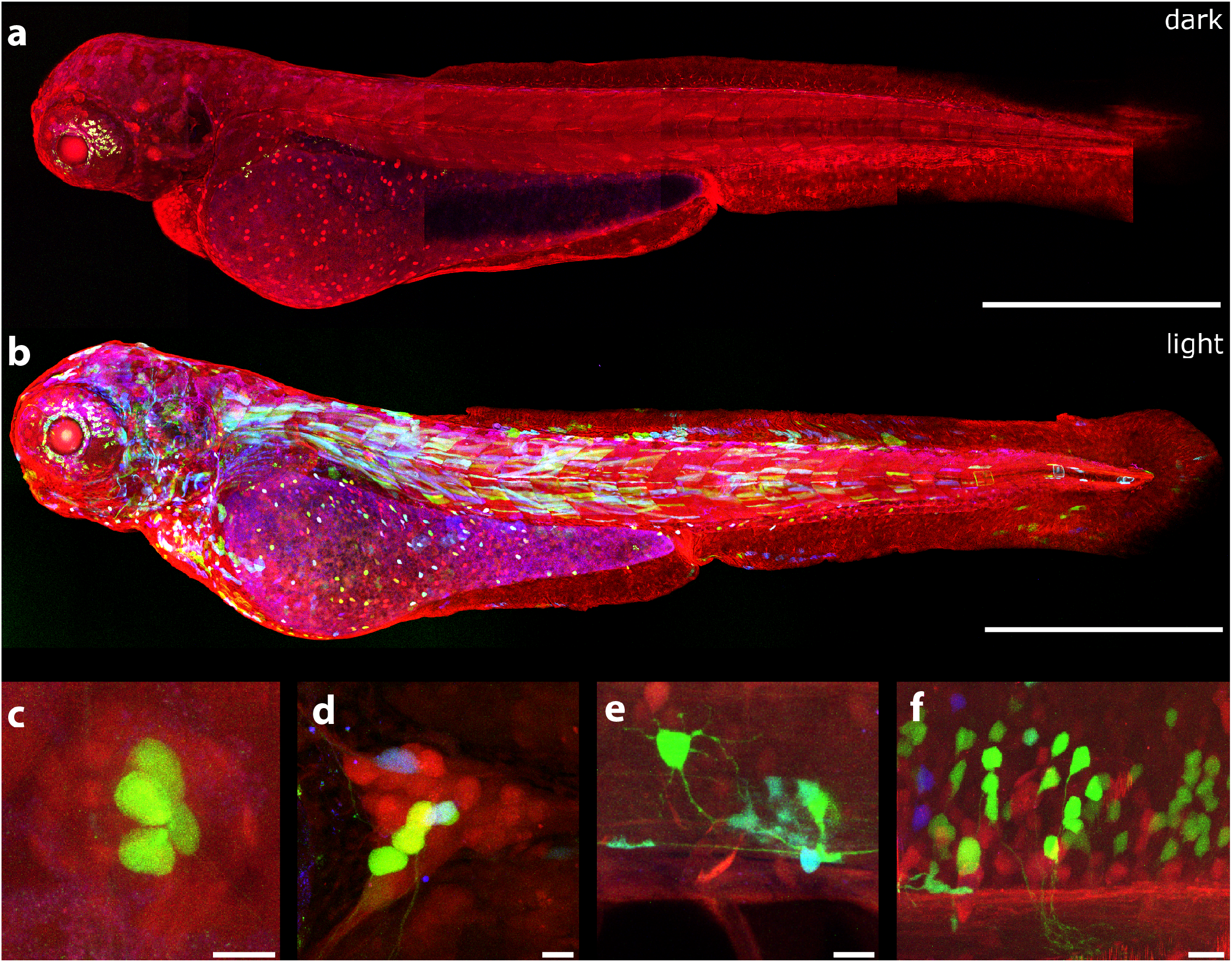
CreV allows tight optogenomic modifications in multiple tissues of Danio rerio. (**a**) Confocal image of dark reared *Tg(ubi:Zebrabrow-M)* zebrafish larva (3dpf) co-injected with *ubi:N-CreV* and *ubi:C-CreV* plasmids - see Methods section for further details. No recombination was observed. In the default state, RFP is expressed in all cells. Apparent green signal is due to autofluorescence. (**b**) Confocal image of a *Tg(ubi:Zebrabow-M)* zebrafish larva (3 dpf) co-injected with *ubi:N-CreV* and *ubi:C-CreV* plasmids and immediately exposed to light. CreV-mediated recombination in multiple tissue types is reflected as expression of YFP and CFP: (**c**) lateral line hair cells, (**d**) trigeminal ganglion, (**e**) spinal neuron and glial cells, and (**f**) hindbrain neurons. Scale bars (**a-b**) 500μm, (**c-f**) 10μm.

We next compared iCreV and iDreV, with Magnet based versions. In cell culture Magnet based constructs induced significant amount of background recombination in the absence of light. iDreV and iCreV exhibited control levels of recombination in the absence of light and retained high levels of inducibility – ~1.6 fold higher for iCreV as compared to split CreV and ~0.6 fold lower for iDreV compared to split DreV in relative fluorescence intensities (**Supp. Fig. 3b)**. For both iCreV and Cre-Magnets efficient recombination was observed *in vivo* after light stimulation -please see the details of the *in vivo* experiments below. However, in no light control cases with Cre-Magnets, background recombination was observed throughout the brain structures − ~50 cells per 100 μm coronal slice, whereas iCreV did not lead to significant recombination (**Supp. Fig. 4**).

**Figure 4.**
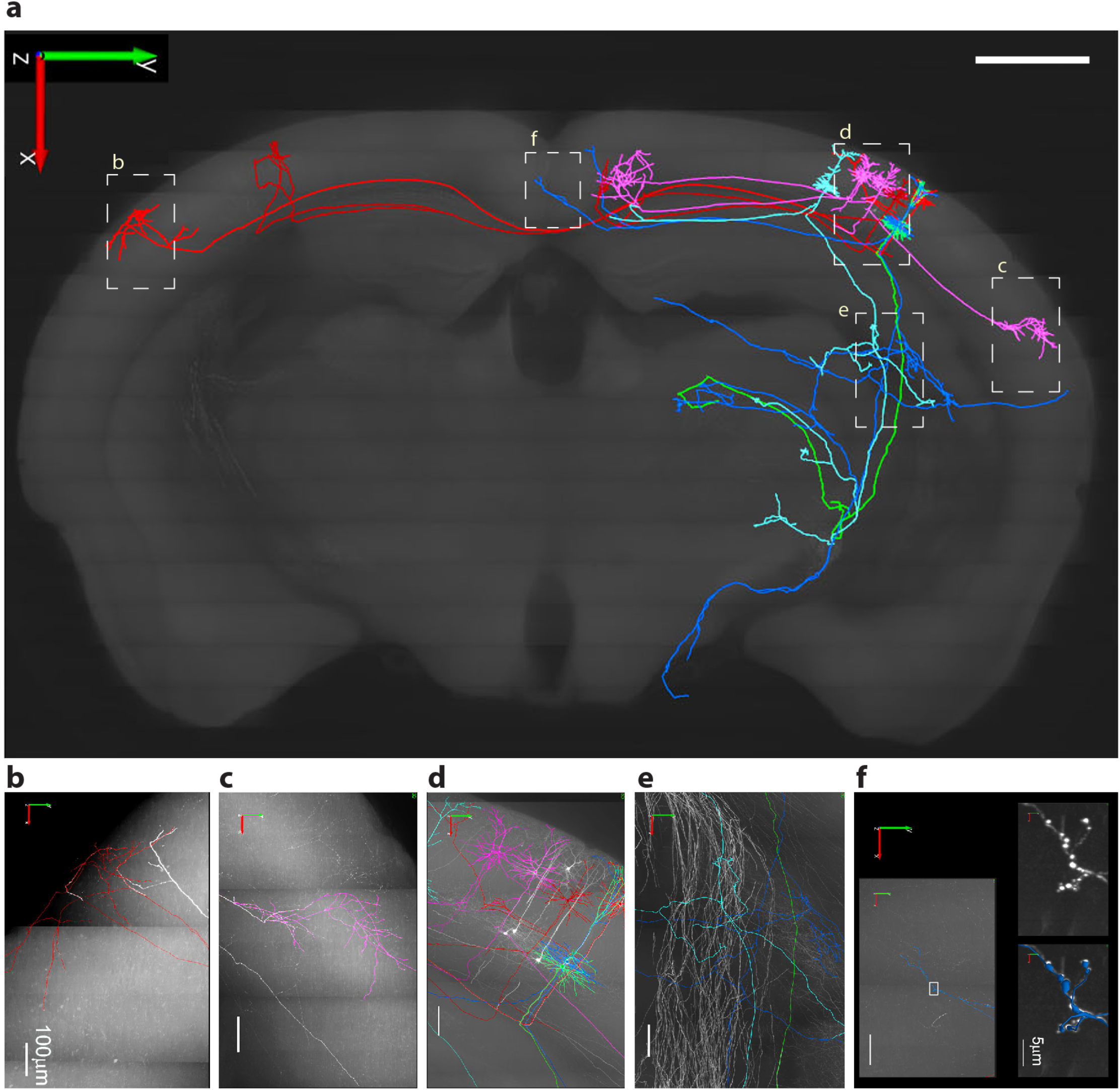
Cortical pyramidal cells (PCs) labeled with CreV and reconstructed at the whole-brain level. (**a**) Reconstructed 8 PCs in mouse somatosensory cortex include 3 layer 2/3 PCs (in pink) with ipsilateral cortico-cortical projections, 2 layer 2/3 PCs (in red) with contralateral cortico-cortical projections, and 3 layer 5 thick tufted PCs (1 in green, 1 in blue, 1 in light blue) with ipsilateral cortico-subcortical projections. Local axonal clusters are incomplete because the labeling at the somata region is still too dense in this brain for tracing of fine axonal branches. The 8 reconstructed PCs are superimposed onto a coronal brain plane located 5201-5400 μm posterior to the olfactory bulb. Scale bar: 1 mm. (**b-f**) Enlarged views of areas outlined by dashed boxes in **a**, with reconstructions (in colors) superimposed on original images with GFP fluorescence in white. In **f**, the two panels on the right (without or with reconstructions in blue) are enlarged views of the boxed area in the left panel, showing the high-resolution details of a segment of axon with enlarged boutons clearly visible. The whole brain image stack (composed of 12089 images, resolution of XYZ, 0.3 x 0.3 x 1 μm) was obtained using fluorescence micro-optical sectioning tomography system (fMOST). 3D reconstruction was performed manually with Neurolucida 360 (NL360).

We also compared the cryptochrome-based light inducible Cre recombination system -CRY2 ^32^, with iCreV. The comparison in cell culture conditions revealed that iCreV performs ~4 fold better after 60 minutes of light stimulation than the CRY2 system, whereas the CRY2 based Cre recombination system allows only a slight increase over baseline recombination (~1.2 fold) (**Supp. Fig. 3c**). We were not able to test the Cry2 based constructs *in vivo* since these genes were too large to fit into the rAAV viral vectors we used.

Our results suggest that under the assayed conditions, VVD-based optogenomic control is tighter and more suitable for certain downstream applications compared to the previously described systems.

### Design and screening of light-inducible Flp recombinases

To increase intersectional genomic modification possibilities, we turned into a third widely used site-specific DNA recombinase: Flp^45^. There isn’t enough protein sequence similarity between Flp, a recombinase from yeast, and Cre or Dre, recombinases from bacterial phages to guide a split based on homology. Thus, we resorted to structure-based *de novo* design and screening for split sites based on crystal structure of the Holliday junction bound Flp^46^. Using a more efficient, codon-optimized variant of Flp (FlpO)^47^; we made Flp splits at 21 loop locations that correspond to transitions between alpha helices and/or beta sheets with hopes to not alter overall functionality within the dimer form. We generated these VVD-fused split iFlpV constructs using the iCreV backbone by replacing the N- and C-termini of Cre with the 21 split Flp variants (**Supp. Fig. 5a**).

**Figure 5.**
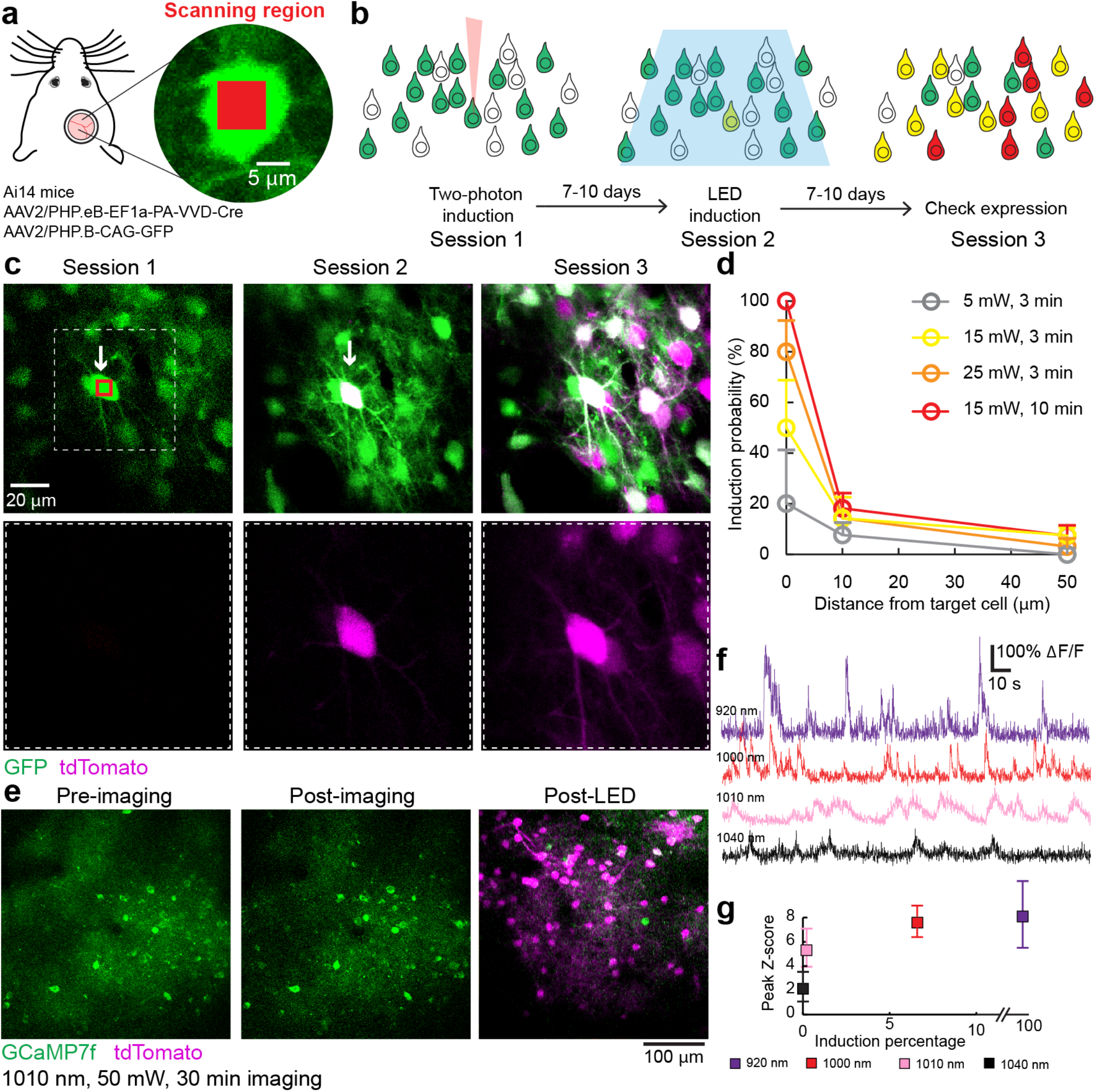
Two-photon guided targeted single cell optogenomic modifications by iCreV in mouse neocortex. (**a**) Diagram of the mouse preparation for the *in vivo* experiment. The insert shows laser induction area (red square) in the target cell (green) within the mouse’s craniotomy. Induction scanning was performed by a Galvo-Galvo scanner. Viral vectors and mouse line used are listed (see Methods section for details). (**b**) Experimental timeline. Each mouse underwent three imaging sessions with two-photon targeted induction (session 1), LED induction (session 2) and final assessment for expression (session 3). (**c**) Example 2P images from a targeted induction experiment. Arrows indicate the target cell and a red box indicates the induction scanning region. Bottom row shows zoomed in regions (dashed box) containing the target cell. (**d**) Quantification of the *in vivo* two-photon induction rate under different scanning protocols at 0, 10, and 50 μm distance to target cells. N = 3-5 cells for target cells, 13-22 cells for 10 μm distance, and 85-122 cells for 50 μm distance. Error bars represent Standard Error of the proportions. (**e**) Example two-photon images before, and after 30 minutes of calcium imaging with 1000-nm two-photon excitation of jGCaMP7f. (**f**) Example calcium traces from the imaged cortex at different wavelengths. GCaMP signal was collected at 30 Hz with a resonance scanner. (**g**) Quantification of the *in vivo* two-photon induction rate after 30 minutes of calcium imaging under four excitation wavelengths held at 50 mW, and the quality of calcium signal in each condition. Data shown as median with error bars indicated for 25 and 75 percentile of calcium peak height. N = 100 events for all conditions.

After gene synthesis, these constructs were cloned into CMV promoter-driven mammalian expression plasmids and tested for light-induced recombination in cell culture via transient transfections together with a Flp-dependent fluorescent reporter plasmid. Among the variants, iFlpV2, 19 and 20 gave the most significant light-induced recombination (**Supp. Fig. 5b**). Dark conditions for all these cases were almost identical to the controls indicating that, like for iCreV and iDreV, the background recombination was almost non-existent.

In an effort to further improve the efficiency of light-inducible Flpase activity after this initial screen, we designed 62 more iFlpV variants to scan Flp split sites surrounding those of iFlpV2, 19 and 20 (**Supp. Fig. 5c**). Among all these constructs FlpV2 still yielded the highest efficiency.

### Light-inducible recombination with RecVs with whole-brain infections in vivo

Recently, a new blood brain barrier permeable rAAV serotype, PHP.eB, was reported to be effectively infecting large proportions of the entire nervous system in mice after retroorbital delivery^48^. Thus, we reasoned this technique could be a good and rapid approach for examining RecV constructs in the entire mouse brain. We examined background recombination in the absence of light in any part of the brain, as well as spatial restriction of light inducible recombination.

For this purpose, we generated an EF1α promoter-driven Cre-expressing rAAV as well as RecV rAAVs of the PHP.eB serotype. We found that the PHP.eB EF1a-Cre virus, when injected either intracerebroventricularly (ICV) or retroorbitally into the fluorescence Cre-reporter mice, efficiently infects the entire brain relatively homogenously (**Fig. 2a, Supp. Fig. 6a**). To our knowledge, this is the first demonstration of effective PHP.eB mediated gene delivery into the brain using the ICV route. This method may help to further restrict the infection to the nervous system and may help overcome obstacles related to intravenous delivery of genes for both basic research as well as gene therapy field applications.

We next tested the light-inducible recombination using RecVs with whole brain infections. To do so first we injected a mixture of PHP.eB NDreV and CDreV viruses into the right ventricle of the Dre-dependent fluorescent reporter mouse, Ai66R. After an incubation period of two weeks, the left hemisphere was exposed to light. A gradient of recombination could be observed from the top of the left hemisphere to deeper structures (**Fig. 2b**). Deeper brain structures and sites far away from the light stimulation did not have any fluorescently labeled cells-except for ICV needle tract where recombination took place likely due to multiple copies of infection. In addition, we tested the iCreV construct with PHP.eB serotype, delivered by ICV or RO routes, in a Cre-dependent nuclear reporter mouse line, Ai75. Very similarly, we observed substantial recombination in the hemisphere exposed to light regardless of the viral delivery route (**Fig. 2c and Supp. Fig. 6b**). Equivalent tests of the iDreV construct in a Dre reporter line yielded a gradient of recombination from the light stimulated hemisphere (**Fig. 2d**). As an additional control iCreV-injected Cre-dependent reporter mice were maintained without light stimulation for four weeks after initial injection. No significant amount of recombination was observed, indicating tight optogenomic control under these conditions (**Supp. Fig. 6d**). Finally, equivalent tests of the iFlpV construct in the Flp reporter mouse line resulted in efficient and specific light-induced recombination (**Fig. 2e**). These results indicate that the technique allows highly specific light-inducible recombination.

Thus far all the recombinase reporter mouse lines we used were generated via targeted insertions into the *Rosa26* locus^49^. To test if a different locus could be modified by the RecV system, we RO injected PHP.eB iCreV rAAVs into the Ai167 ChrimsonR reporter mouse line, which is inserted into the Tigre locus^50^. We again observed efficient recombination shaped as a gradient from the light-stimulated left hemisphere confirming that other genomic loci in mice can be modified by the RecV system (**Supp. Fig. 6c**).

We also tested the feasibility of light inducible recombination within a deeper brain area, striatum. Deep brain imaging or recording is more challenging due to tissue damage and blood clotting; both create obstacles for light passing through the tissue. By utilizing 1P-light through an optical fiber we were able to induce local recombination via iCreV in the striatum of Ai162 GCaMP6s Cre reporter mice^50^, and record changes in fluorescence due to calcium concentration dynamics before and after light stimulation (**Supp. Fig. 7a-d**). Due to the localized illumination, GCaMP6s was expressed directly under the fiber, in comparison to the wide expression of red fluorescence coming from the control virus (**Supp. Fig. 7b**). This expression pattern is beneficial for calcium imaging and recording, as it allows expression directly at the desired location. Our results show that light-mediated genomic modifications can be efficiently spatiotemporally regulated *in vivo* also within deep brain targets therefore providing a tool for restricted reporter expression under the optical device.

### Cell-class specific targeting by intersection of viral RecVs and transgenic recombinases

Versatile and refined cell type targeting can be achieved by combined use of two or more recombinases with distinct recombination activities^50^. For example, new RecV tools could be combined with existing transgenic recombinase lines and intersectional reporters.

To test the feasibility of this approach, we co-injected PHP.eB iFlpV viruses together with a PHP.eB Cre/Flp dependent fluorescent reporter virus into the cortex of mice containing the Rbp4-Cre-KL-100 transgene, which drives Cre expression mostly in L5 excitatory cortical cells. After unilateral light stimulation we observed layer 5 specific reporter gene expression within the light-stimulated hemisphere in these mice (**Fig. 2f**), suggesting high degree of intersectional specificity under the tested conditions.

To provide further evidence that light-mediated intersectional targeting can be achieved in other neuron types, we used the Sst-IRES-FlpO mouse line, which expresses FlpO recombinase selectively in inhibitory somatostatin (Sst) neurons, in conjunction with iCreV. We crossed this mouse line to a Cre/Flp double dependent fluorescent reporter mouse line Ai65. The resulting double-positive mice were retro-orbitally injected with PHP.eB iCreV viruses to infect the entire brain relatively homogenously. Reporter expression was observed in sparsely distributed neurons close to the light stimulation site across multiple cortical layers (**Fig. 2g**). To ascertain that the recombination we observed within the light induced hemisphere is indeed specific to the Sst-positive neurons, we performed immunohistochemistry and found that all the fluorescently labeled neurons were also immune-reactive to the SST antibody (**Fig. 2g**). These results confirm that tight intersectional control can be achieved by combining virally delivered iFlpV or iCreV with transgenic class-specific Cre or FlpO, respectively.

### CreV induces tight light-dependent recombination in zebrafish embryos in many tissue types

To determine if RecVs could effectively induce light-dependent recombination in other loci and organisms other than mice we tested it in another model organism: Danio rerio (zebrafish). We co-injected plasmids containing either N-CreV or C-CreV under the control of the ubiquitous ubiquitin (*ubi*) promoter into single cell zebrafish embryos of transgenic zebrabow strain tg (*ubi:Zebrabow*)^51^ and reared embryos either in the light or the dark for 72 hours. In this line within the default state, RFP is expressed in all cells. After Cre-dependent recombination, YFP and/or CFP are expected to be expressed. Of the 24 viable embryos reared in the dark 1/24 displayed any recombination as measured by the presence of YFP or CFP. Of the 37 viable embryos reared in the light nearly all (32/37) had YFP or CFP expression, confirming that CreV-induced recombination was light dependent. We observed recombination in many tissues including muscle, skin, heart, spinal cord neurons and non-neuronal cells, hindbrain, trigeminal ganglion, Rohon-Beard sensory neurons and hair cells of the lateral line (**Fig. 3a-f**), suggesting that CreV is effective at inducing recombination across many cell types and tissues.

### RecV-mediated sparse labeling enables whole-brain reconstruction of single neuron morphologies

The development and *in vivo* validation of the light-inducible recombinase system (RecV) opens doors to many applications. Here we present several of such applications, mainly using light to control the precise location and number of neurons labeled to achieve the desired sparsity of targeted neuronal populations for their visualization, functional imaging, as well as reconstruction of single neuron morphologies across the entire brain to understand their axonal projection specificity.

Cre-mediated *in vivo* reporter expression using a Cre reporter mouse line Ai139 results in strong expression of GFP in multiple cortical layers. Ai139 is a TIGRE2.0-reporter line expressing very high level of GFP (via tTA-mediated transcriptional amplification) allowing visualization of detailed cellular morphologies^50^. We tested if light applied to the tissue could sparsely induce CreV, to result in sparse yet strong expression of the reporter at the single cell level, which would enable whole-brain reconstruction of single neuron morphologies.

For this purpose, we injected the cortex of Ai139 mice with the 1:1 mixture of NCreV and CCreV rAAVs in serial dilutions and applied various durations of light across the skull to induce recombination in a sparse population of cortical neurons. We then imaged the whole brains using the fluorescence micro-optical sectioning tomography (fMOST) technique^52^. Lower dose of CreV viruses and shorter duration of light induction led to sparse and strong labeling of individual neurons (**Supp. Fig. 8**). Sparseness was low enough that axons from many neurons could be traced. In the example brain shown in **Figure 4** (also see **Supp. Movies 1 and 2**), 8 primary somatosensory cortical neurons were manually reconstructed. These neurons include 3 Layer 2/3 pyramidal cells (PCs) with ipsilateral cortico-cortical projections, 2 Layer 2/3 PCs with contralateral cortico-cortical projections, and 3 Layer 5 thick tufted PCs with ipsilateral cortico-subcortical projections, revealing distinct axonal projection patterns.

In many cases where restricted genetic access is not possible with conventional methods, it is desirable to further confine the region of recombination by precise induction through localized illumination. To demonstrate this ability, we performed headpost craniotomy and two-photon (2P) assisted local laser stimulation experiments in trained head-fixed Ai139 reporter mice. To achieve sparseness, we co-injected a 1:5 mixture of rAAV iCreV and rAAV tdTomato-expressing viruses into the primary visual cortex. The tdTomato virus was included to guide our light stimulation, as well as to serve as a control of overall infection breadth. Our results indicate that local and sparse labeling of neurons is achievable using this method. Furthermore, due to the strength of the Ai139 reporter expression, individual axons and boutons can be readily visualized without immunohistochemical enhancement at sites far away from neuronal cell bodies, suggesting that these brains could be subjected to whole-neuron reconstruction (**Supp. Fig. 9**).

### RecVs enable 2P-mediated single cell specific targeted optogenomic modifications in combination with functional imaging in vivo

A powerful application for the light-inducible iCreV system is to identify -e.g. functionally or genetically- individual cells of interests and then activate the Cre-dependent gene expression in these defined cells at will. However, 1P illumination is not suitable for this application, since it inevitably activates the iCreV in cells within the light path in addition to the target. One approach to achieve single-cell specific targeting is through 2P activation of iCreV which minimizes unspecific targeting. To investigate the feasibility of this approach *in vivo* at single cell level, we next tested the iCreV system using 2P induction in mouse somatosensory cortex (**Fig. 5a-b**).

We co-injected iCreV and GFP-expressing viruses in the somatosensory cortices of a Cre-dependent red fluorescent reporter mouse line (Ai14). The GFP fluorescence served as a landmark for target cells. After identifying the target cell, we focused the laser in the middle of the cell soma (potentially nuclear region), and then laser-scanned within this small region - as a window of ~6×8 μm- under various conditions of laser power and time. Seven to ten days after this initial induction session, we re-imaged the target cells to examine the expression of tdTomato. In order to reveal all the inducible cells in the region, we then exposed the neurons to 1P LED light for 30 minutes, and then counted all the tdTomato-expressing cells seven days later.

We showed that 2P induction of iCreV led to Cre-dependent gene expression in target cell with single-cell precision. In many cases, we observed potent tdTomato expression in the target cell but not in the inducible cell right next to it (**Fig. 5c and Supp. Fig. 10**), demonstrating the high spatial specificity. We tested various 2P induction protocols and plotted the probability of iCreV activation against the distance to the target cell in each condition (**Fig. 5d**). The results showed that target cell induction rate increased as more laser power and scan time was applied during the induction. However, powerful induction in some cases also led to nonspecific induction within 10 μm radius of the target cell (as high as 18%), which may result from the activation of the iCreV constructs in the neural processes surrounding the target cell, or from micron-level movement of the mouse brain during the induction window. Cells within the 10-50 μm radius from the target cell showed a baseline induction rate of 6-7%. Since the induction rate observed in session 1 was very low (~3% on average, **Supp. Fig. 11**), this non-targeted induction was likely due to the non-specific activation of iCreV by ambient light during mouse handling, by the two-photon stimulation during acquiring the Z-stack images or by spurious recombination due to very high multiplicity of infection with the viral vectors.

We next tested the feasibility of combining calcium imaging with the iCreV system, by measuring the induction probability of iCreV after 30 minutes of calcium imaging. We co-injected iCreV and GCaMP7f^53^ viral constructs into Ai14 mice cortex. 920 nm excitation provided good quality jGCaMP7f signal, however, was also potent in activating the iCreV system (close to 100%). To reduce iCreV induction during the imaging session, we tested longer wavelength excitation of jGCaMP7f at 1000, 1010 and 1040 nm. At these wavelengths, the induction of iCreV was significantly reduced, with moderately reduced jGCaMP7f signals (**Fig. 5e-g**).

## DISCUSSION

In this study, we used wild-type VVD to create three light-inducible recombinases Cre, Dre, and Flp, thus expanding the repertoire of optogenomic manipulation tools even further. We also showed that these light-inducible recombinases work highly efficiently and intersectionally in the mouse brain to label specific cell classes or types. We further demonstrated that RecVs allow effective light induced optogenomic modifications in multiple loci within the mouse genome and zebrafish.

We provided proof-of-principle experiments showing that RecV mediated light-inducible site-specific DNA modifications are possible in the mammalian nervous system at single cell level. We provided examples of single-cell soma-targeted 2P-mediated optogenomic modifications and established imaging and conversion parameters to induce such modifications with unprecedented spatial resolution. Our data also provide a quantitative description of the induction specificity of iCreV at different 2P wavelengths and support the feasibility of combining calcium imaging with iCreV system in mice *in vivo*.

Light-inducible Cre recombinases were reported previously^32,33,54^, and they were mainly based on two types of light-inducible protein dimerization systems: CRY2-CIB1 and Magnets. A direct comparison of an improved CRY2 based ^32^ and the VVD based system presented in this study shows that the CreV system may be preferable due to its higher level of inducibility under the conditions tested *in vitro*. Additionally, our results indicate that the heterodimeric Magnet-based light-inducible Cre recombination is efficient, but it is also relatively leaky compared to our VVD-based counterpart both *in vitro* and *in vivo*. By incorporating the improved designs reported in the Magnet system^33^ we were able to produce efficient RecV co-expression constructs, thereby decreasing the number of independent components that need to be delivered to the experimental animal.

The demonstration of the ICV route for efficient whole-brain infection using PHP.eB AAV viruses could be valuable in many scenarios. For example, it could infect the brains of embryos in which RO vein is too small to inject, or to avoid damage to the eye of the embryo since in mice RO route requires penetrating a needle next to the eye. This approach may further help focus infection to the nervous system and overcome obstacles related to intravenous delivery of genes for basic research in a multitude of mammalian species. Genetic modifications of larger animal models such as primates are expensive. Intravenous delivery in the adult primate may require substantial number of viral particles. Thus, if the PHP.eB viruses are delivered by the ICV route during adulthood or development, it may help infect majority of the nervous system with much less virus. This approach may also overcome immune-response related issues that interfere with infections. Finally, this method may be useful in clinical applications for gene therapy purposes.

The spatiotemporally selective targeting of cell populations or individual cells using light could allow fine-scale combinatorial functional, anatomical and circuit-level studies of cell type-specific networks. In addition to combining opsin expression with iCreV, and achieving efficient GCaMP expression using iCreV, the RecV viral vectors and Cre/Flp-dependent iDreV or iFlpV mouse lines can be combined with existing cell type-specific genetic tools, and RecVs can also be used to generate new driver lines. New RecV versions incorporating drug-inducible (e.g. tamoxifen^55^ or trimethoprim (TMP) inducible^56^) recombinases can be developed to further reduce background recombination and enhance temporal specificity. With this approach, one can envision performing a variety of combinatorial experiments including single-cell activity imaging using genetically encoded voltage^57,58^ or calcium indicators^59^; single cell monosynaptic rabies tracing^6,7^ or targeted optogenetic and chemogenetic protein expression^60^, all in a noninvasive or minimally invasive manner. The spectral and temporal separation as well as drug-inducible versions could enable sequential investigations of functionally relevant cell populations, followed by light mediated DNA recombination to selectively activate effector gene expression in specific cells of interest, to examine their structure and connectivity and/or to perturb their function.

This technology could also enable loss/gain-of-function studies by switching genes off or on with much refined regional and cell type specificity, followed by monitoring of effects on physiology or animal behavior. One could also employ this method to study development by switching genes on or off with high temporal precision during the rapid cascade of events throughout the proliferation and differentiation processes. The half-life of the VVD homodimer is ~2 hours after brief light activation^13^. One can try to apply targeted light at very precise time points using standard or gentler *in utero* techniques. This may be advantageous over the current CreER-based genetic fate mapping studies^61^ as it would eliminate the tamoxifen side effects as well as provide spatially and temporally more precise recombination. Furthermore, this approach could allow highly targeted neuronal network reconstructions by selectively labeling neurons in particular locations or having particular activity patterns or functional properties (*e.g.*, using 2P induction). Sparse expression of reporter genes using diluted Cre viruses or low doses of tamoxifen in CreER mice already exist. However, with RecVs one can perform even more spatiotemporally restricted induction, and it can be combined with prior functional characterization using 2P imaging and optical indicators in an intersectional manner.

The RecV approach can also be used in other species of interest in which light could be used to access individual cells of interest. For model systems in which germline genetic modification is feasible, the experiments can be designed to integrate multiple components of the described systems into the germline^62^. In model systems in which germline modification is not available or is prohibitively expensive, these experiments can be performed by using viral vectors in which a certain level of cell-type specificity may be achieved by using short cell type specific promoters or enhancers^63,64^, or target-defined specificity may be achieved with highly efficient retrogradely infecting designer viruses^65^.

Overall, the broad range of potential applications show that the light-inducible recombinase system should enable much improved spatiotemporal precision and multiple combinatorial strategies for the micro- and macro-level analyses of neural circuits as well as many other biological systems in a multitude of organisms.

## METHODS

### Plasmid and virus construction for RecVs

Sequences of NCre-5G-VVD, VVD-5G-CCre, NCre-5G-VVD-IRES-VVD-5G-CCre, VVD-5G-CCre-IRES-NCre-5G-VVD, NCre-5G-VVD-PQR-VVD-5G-CCre, VVD-5G-CCre-PQR-NCre-5G-VVD, VVD-5G-CDre-IRES-NDre-5G-VVD, NCre-Magnets-NLS-P2A-NLS-Magnets-CCre, NCre-VVD-NLS-P2A-NLS-VVD-CCre, NDre-Magnets-NLS-P2A-NLS-Magnets-CDre, NDre-VVD-NLS-P2A-NLS-VVD-CDre, and all iFlpV versions were chemically synthesized (GenScript, Piscataway, NJ). 19-59 amino acid N terminus and 60-343 C terminus of Cre were used in all CreV cases. To screen poly-cistronic cassettes with the best light-inducible recombinase activity, N and C parts of RecV, as well as IRES-mediated, PQR-mediated and P2A-mediated RecV poly-cistronic cassettes were cloned into pCDNA3.1 with the CMV promoter (**Supp. Fig. 1**).

To generate recombinant AAV viruses expressing split VVD-Cre (CreV) or VVD-Dre (DreV), the N or C part of CreV and DreV were cloned after the human EF1a promoter, followed by WPRE and hGH-polyA signal (**Supp. Fig. 1**). The Cre reporters, pAAV-EF1a-Flex-dTomato or EGFP-WPRE-hGHpA, used pairs of double inverted LoxP and Lox2272 sites to flank the reporter dTomato or EGFP sequence. The Dre reporter, pAAV-EF1a-Frex-dTomato-WPRE-hGHpA, was generated by inserting an inverted dTomato sequence flanked with Rox sites after the human EF1a promoter, followed by WPRE and hGH-polyA signal (**Supp. Fig. 1**).

21 iFlpV variants were generated with custom gene synthesis as follows: iFlpV1: 11 amino acids (aa) N and 412 aa C; iFlpV2: 27 aa N and 396 aa C; iFlpV3: 49 aa N and 374 aa C; iFlpV4: 67 aa N and 356 aa C; iFlpV5 72 aa N and 351 aa C; iFlpV6: 85 aa N and 338 aa C; iFlpV7: 95 aa N and 328 aa C; iFlpV8: 114 aa N and 309 aa C; iFlpV9: 129 aa N and 294 aa C; iFlpV10: 151 aa N and 272 aa C; iFlpV11: 169 aa N and 254 aa C; iFlpV12: 197 aa N and 226 aa C; iFlpV13: 208 aa N and 215 aa C; iFlpV14: 237 aa N and 186 aa C; iFlpV15: 251 aa N and 172 aa C; iFlpV16: 290 aa N and 133 aa C; iFlpV17: 318 aa N and 105 aa C; iFlpV18: 343 aa N and 80 aa C; iFlpV19: 374 aa N and 49 aa C; iFlpV20: 388 aa N and 35 aa C; iFlpV21: 408 aa N and 15 aa C. Additional iFlpV2 variants were generated spanning amino acids 16-39 and 366-405 covering the entire region leading to 61 additional constructs. Construct 62 was generated based on iFlpV2 with an addition of the linker GGSGG present between the C terminus VVD and FlpV to also N terminus FlpV and VVD. These constructs were cloned in pcDNA3.1 mammalian expression plasmids.

AAV1, AAV DJ and AAV PHP.eB serotype viruses were produced in house with titers of AAV1-EF1a-NCreV, 1.05 × 10^12^ genome copies (GC); AAV1-EF1a-CCreV, 5.16 × 10^12^; AAV1-EF1a-NDreV, 4.20 × 10^13^; AAV1-EF1a-CDreV, 5.40 × 10^13^; AAV-DJ-EF1a-Cre, 2.00 × 10^13^; AAV1-CAG-Flex-EGFP, 1.34 × 10^13^; AAV-DJ-EF1a-Frex-dTomato, 1.90 × 10^12^; 7.7 × 10^11^; 1.6 × 10^13^; AAV-PHP.eB-EF1a-Cre, 5.8 x10^13^; AAV-PHP.eB-Syn-NDreV, 4.2 × 10^13^; AAV-PHP.eB-EF1a-CDreV, 3.9 × 10^13^; PHP.eB iCreV, 2.6 × 10^13^; PHP.eB iDreV, 3.3 × 10^13^, PHP.eB iFlpV, 2.7 × 10^13^, AAV-PHP.eB-EF1a eGFP 2.03E+13 and AAV- Cre-Magnets 3.00E+13 per ml. AAV5.CAG.tdTomato (1.0 × 10^13^ GC/mL) were purchased from UNC Vector core.

### Light activation in cultured cells

HEK293T cells were seeded into 6-well plates one day before transfection and reached 80% confluency on the day of transfection. Cells were co-transfected with a reporter expressing dTomato –for Cre and Flp- or dTomato-for Dre- for Cre, Dre or Flp mediated recombination and various constructs of split RecVs. Cells in the control groups were transfected with reporters alone. Each condition contained 4 replicates. Plates were kept in dark immediately after transfection. Twenty-four hours later, cells were exposed to blue light, and were then kept in dark immediately after light exposure. Cells were imaged for fluorescent reporter expression 48 hours after light induction, using an inverted fluorescence microscope. RecV activated dTomato expression in each condition was quantified using Image J.

### Surface/Cortical In vivo 1P optogenomic modifications

Stereotaxic injections were made into adult C57BL/6J (stock no. 00064, The Jackson Laboratory, Bar Harbour, ME) or transgenic reporter mice with a 1:1:1 or 1:1 mixture of three or two different rAAVs. For all experiments, animals were anesthetized with isoflurane (5% induction, 1.5% maintenance) and placed on a stereotaxic frame (model no. 1900, David Kopf Instruments, Tujunga, CA). An incision was made to expose the skull, including bregma, lambda, and the target sites. Stereotaxic coordinates were measured from Bregma and were based on The Mouse Brain in Stereotaxic Coordinates^66,67^. A hole was made above the target by thinning the skull using a small drill bit until only a very thin layer remained. An opening was then made using a microprobe, and the remaining thinned skull was gently pulled away. All animals were injected at each target with 500 nl of virus at a rate of ~150 nl/min using a Nanoject II microinjector (Drummond Scientific, Broomall, PA). Intraventricular injection of AAV-PHP.eB viruses was conducted by injecting 2 μl of virus into the lateral ventricle using a Nanoject II microinjectior. The glass pipettes had inner diameters between 10-20 μm.

Two weeks following AAV injection, animals were anesthetized and returned to the stereotaxic frame. An incision was made in the previous location to once again reveal the location of the injection sites. An LED light source (LED-64s, Amscope, Irvine, CA) was mounted to the surgical microscope and positioned 3-4 inches directly above the animal’s skull. The amount of time the animal was exposed to light was varied by experiments. Small amounts of sterile PBS were periodically applied to the scalp and skull to prevent drying.

Two weeks following light exposure, animals were perfused with 4% paraformaldehyde (PFA). Brains were dissected and post-fixed in 4% PFA at room temperature for 3–6 h and then overnight at 4°C. Brains were then rinsed briefly with PBS and stored in PBS with 10% sucrose solution. Brains were then sectioned at a thickness of 100 μm while frozen on a sliding microtome (Leica SM2010 R, Nussloch, Germany). Brain sections were mounted on 1×3 in. Plus slides and coverslipped with Vectashield with DAPI (H-1500, Vector Laboratories, Burlingame, CA). Slides were then imaged using a 10x objective on a Leica TCS SP8 confocal microscope (Leica Microsystems, Buffalo Grove, IL).

For fMOST imaging, two weeks or longer following light exposure, animals were perfused with 4% paraformaldehyde (PFA). Brains were dissected and post-fixed in 4% PFA at room temperature for 3–6 h and then overnight at 4°C. Brains were then transferred to PBS with 0.1% sodium azide for storage at 4°C until embedding.

### Surface/Cortical in vivo population 2P optogenomic modifications

A titanium head plate was attached to the skull of mice to allow positioning and restraint of the animal during imaging. The hole of the head plate was positioned over visual cortical areas, approximately 2.9 mm posterior and 2.7 mm lateral from Bregma. A 5 mm craniotomy was cut into the skull using a dental drill. The dura was then removed, and a multilayer glass coverslip was positioned above the craniotomy. The head plate and coverslips were secured using cyanoacrylate glue and metabond. After a period of at least one week, a dental drill was used to remove the cement and metabond holding the coverslip in place, and the coverslip was removed. A Dumont Nanoject II was then used to inject 500 nL of viruses into visual cortex. A new coverslip was placed and adhered. The area above the coverslip was blocked from light using a combination of dental cement and Kwik-cast, both mixed with black acrylic paint powder.

After at least 3 weeks following viral injection, the animal received two-photon laser stimulation. Under dark conditions, the Kwik-cast was removed, and the animal’s head plate was mounted in position. The injection area was identified by the presence of the EGFP labelled cells. Laser output was set to 900 nm to optimally induce recombination. A 600 × 600 μm area was stimulated at three depths (100 μm, 150 μm, and 200 μm) for 15 minutes each. After stimulation, black Kwik-cast was reapplied. Two weeks following stimulation, mice were perfused.

### Deep brain in vivo stimulation/imaging experiments

For deep brain optogenomic modification and imaging experiments stereotaxic injections were made into Ai162-GC6s (Stock No. 031562, The Jackson Laboratory, Bar Harbour, ME) Cre dependent GCaMP6s reporter mice with a 1:1 mixture of PHP.eB.iCreV and a control AAV unconditionally expressing red fluorescent protein (AAV5.CAG.tdTomato) into the striatum. For all experiments, animals were anesthetized with isoflurane (5% induction, 1.5% maintenance) and placed on a stereotaxic frame (942, David Kopf Instruments, CA, USA). An incision was made to expose the skull, including bregma, lambda, and the target sites. Stereotaxic coordinates were measured from Bregma and were based on The Mouse Brain Atlas^66,67^. A full hole was made above the target. All animals were injected with 400 nl × 2 of virus mixture, at two dorsoventral positions, 300um apart, at a rate of ~80 nl/min using UltraMicroPump (UMP3-4, World Precision Instrument, Sarasota, FL). Following virus injection, an optical fiber with cut length of 5 mm and diameter of 400 μm (NA 0.48, Doric lenses, Quebec, QC, Canada) for 1P stimulation was firmly mounted to a stereotaxic holder. The optical fiber was then inserted to the striatum (AP +1.0 mm, ML ± 1.3 mm, DV −3.5 mm, from either left or right side) through a craniotomy and positioned 300 μm above the deeper viral injection site. A thin layer of metabond was applied on the skull surface to secure the fiber. In addition, a thick layer of black dental cement was applied to secure fiber implant for 1P illumination to allow positioning and restraint of the animal.

One week following AAV injection of 1:1 virus mixture of PHP.eB.iCreV and AAV5.CAG.tdTomato into GCaMP6s reporter mice, animals’ baseline signals were recorded with a fiberphotometry rig for 10 minutes in the home cage. Fiberphotometry is a method for measuring population calcium dependent fluorescence from genetically defined cell types in deep brain structures using a single optical fiber for both excitation and emission in freely moving mice. Detailed description of the system can be found elsewhere^68^. After recording, mice were connected to a 447nm laser (Opto Engine LLC, UT) using a 200 μm optical fiber, illuminated with 5mW, 100ms pulses, 1Hz for 30 minutes (TTL-controlled by OTPG_4, Doric lenses, Quebec, QC, Canada), in the home-cage. A week following light exposure, fiberphotometry signal was recorded again for 10 minutes. Fiberphotometry peak detection was performed with MATLAB (R2018a), using ‘findpeaks’ function, using a prominence of 2.5. Mice were perfused 4 weeks after illumination.

Animals were perfused with 4% paraformaldehyde (PFA). Brains were dissected and post-fixed in 4% PFA overnight at 4°C. Brains were then rinsed briefly with PBS and then sectioned at a thickness of 100 μm on a vibratome (VT1200 Leica Biosystems, IL, USA). Brain sections were incubated in a blocking solution, containing 1x PBS solution with 0.1% Triton X-100 and 10% normal donkey serum (NDS; Jackson ImmunoResearch, PA, USA) for at least an hour, washed, and further incubated with blocking buffer containing primary antibody (see below for details) at 4C overnight. Afterward, sections were thoroughly washed three times (15 min each) in 1x PBS and then transferred into the blocking solution with secondary antibody (see below for details) for 2h at room temperature. Finally, sections were again washed by 1x PBS solution four times (15 min each), mounted on glass microscope slides (Adhesion Superfrost Plus Glass Slides, #5075-Plus, Brain Research Laboratories, MA, USA), dried, and coverslipped with mounting media (Prolong Diamond, P36965, Thermo-Fischer, CA, USA). For primary antibody: chicken anti-mCherry (1:1000; ab205402, Abcam, Cambridge, UK) was used. For secondary antibody, anti-chicken Alexa Fluor 594 (1:500; 703-585-155, Jackson ImmunoResearch) was used. Fluorescent images from brain tissue were acquired with an LSM 880 confocal microscope (Carl Zeiss, Jena, Germany). We used a 10x Plan Apochromat air objective (NA 0.45), 25x Plan Apochromat water immersion objective (NA 1.2) and three laser wavelengths (488 nm, 561 nm, and 633 nm). Image acquisition was controlled by Zen 2011 software (Zeiss), which also allowed automated tiling, and maximum intensity projection. Images were not further processed. Expression counts were done by summation of the values of the fluorescence within 1mmX1mm below the fiber tip subtracted with the same area at the opposite hemisphere, line by line, and normalized to the maximal value.

### Zebrafish experiments

N-CreV and C-CreV were PCR amplified from pAAV-Ef1a NCreV and pAAV-Ef1a CCreV, and cloned into the pDest-ubi vector (Addgene plasmid # 27323) by Gibson assembly with the following primers: 5’ N-Cre overlap pdest-UBI: attcgacccaagtttgtacaaaaaagcaggctggacgccaccatgacgagtgatgaggtt; 5’ C-Cre overlap pdest-UBI: tcgacccaagtttgtacaaaaaagcaggctggacgccaccatgcatacactgtatgcccc; 3’ polyA (used for both constructs): actgctcccttccctgtccttctgcatcgatgatgatccagacatgataagatacattga

The pDest-ubi:N-vCre-pDest and pDest-ubi:C-vCre mixture containing equal amounts of each plasmid (25pg each) and tol2 transposase RNA (25 pg) was injected into one-cell stage tg(ubi:Zebrabow-M) zebrafish embryos. Embryos were either light or dark reared for 72 hrs. At 3dpf injected embryos were anesthetized with Mesab, mounted in 2% agarose, and imaged on a Zeiss LSM 880 confocal microscope.

### fMOST imaging

All tissue preparation has been described previously^69^. Following fixation, each intact brain was rinsed three times (6 h for two washes and 12 h for the third wash) at 4°C in a 0.01 M PBS solution (Sigma-Aldrich Inc., St. Louis, US). Then the brain was subsequently dehydrated via immersion in a graded series of ethanol mixtures (50%, 70%, and 95% (vol/vol) ethanol solutions in distilled water) and the absolute ethanol solution three times for 2 h each at 4°C. After dehydration, the whole brain was impregnated with Lowicryl HM20 Resin Kits (Electron Microscopy Sciences, cat.no. 14340) by sequential immersions in 50, 75, 100 and 100% embedding medium in ethanol, 2 h each for the first three solutions and 72 h for the final solution. Finally, each whole brain was embedded in a gelatin capsule that had been filled with HM20 and polymerized at 50°C for 24 h.

The whole brain imaging is performed using a fluorescence microscopic optical sectioning tomography (fMOST) system. The basic structure of the imaging system is the combination of a wide-field upright epi-fluorescence microscopy with a mechanic sectioning system. This system runs in a wide-field block-face mode but updated with a new principle to get better image contrast and speed and thus enables high throughput imaging of the fluorescence protein labeled sample (manuscript in preparation). Each time we do a block-face fluorescence imaging across the whole coronal plane (X-Y axes), then remove the top layer (Z axis) by a diamond knife, and then expose next layer, and image again. The thickness of each layer is 1.0 micron. In each layer imaging, we used a strip scanning (X axis) model combined with a montage in Y axis to cover the whole coronal plane^70^. The fluorescence, collected using a microscope objective, passes a bandpass filter and is recorded with a TDI-CCD camera. We repeat these procedures across the whole sample volume to get the required dataset.

The objective used is 40X WI with numerical aperture (NA) 0.8 to provide a designed optical resolution (at 520 nm) of 0.37 μm in XY axes. The imaging gives a sample voxel of 0.30 × 0.30 × 1.0 μm to provide proper resolution to trace the neural process. The voxel size can be varied upon difference Objective. Other imaging parameters for GFP imaging include an excitation wavelength of 488 nm, and emission filter with passing band 510-550 nm.

### Cortical targeted in vivo 2P stimulation of single cells

For *in vivo* targeted single cell optogenomic modifications as well as simultaneous GCaMP7f imaging experiments Stanford Administrative Panel on Laboratory Animal Care (APLAC) approved all animal procedures. Ai14 mice (The Jackson Laboratory, 07908) of two to four months age were used for experiments. For viral vector injection, mice were anesthetized with isoflurane. iCreV virus used: AAV2/PHP.eB-EF1a-iCreV; GFP virus used: AAV2/PHP.B-CAG-GFP; GCaMP virus used: AAV2/9-CamkIIa-jGCaMP7f. Viral vectors were loaded into a glass pipette and injected into cortex with picospritzer (Paker Hannifin). ~500 nL was delivered over 15 minutes and then a 4 mm craniotomy was made with the injection site at the center, 30 minutes after the injection. Dura was removed before cover glass was installed and sealed. A custom-made head bar and cover were secured with dental cement on mouse skull. Imaging experiments started one month after the surgery to allow gene expression.

Mouse was mounted on a running wheel with head fixation and was remained awake during the whole experiment. Before imaging, the head mount cover was removed. In order to operate the mouse and the microscope, a red LED light was used for illumination (WAYLLSHINE). Ultrasound gel (Parker, Aquasonic) was put on the cover glass for the water immersion objective lens. The mouse was aligned manually without checking the focal plane with eye piece to avoid iCreV induction during this process.

For induction experiment, a femtosecond Ti:sapphire laser (Spectra-Physics, Mai Tai) was tuned to 920 nm wavelength. The scanning and image acquisition were achieved with a Prairie (Bruker) two-photon microscope, through a 20X 0.95 N.A. water immersion lens (Olympus XLUMPLFLN-W 0.95 NA 20×). For all the imaging sessions, laser power at a specimen was kept at 25 mW and it was monitored (Thorlabs, PM100D and S130C) at an additional output of the optical path before entering the microscope, whose splitting ratio was calibrated prior the measurements. During the first imaging session, a starting point with unique vessel pattern was identified and recorded as the starting position. All target cells’ relative coordinates to the starting position were recorded and used for relocation in later sessions. Images were acquired with 4 μs pixel dwell time at 1024 × 1024 pixel frame size (field of view 450 micron × 450 micron), with 3 μm Z-axis step size. After identifying a target cell, an induction scanning was carried out using the ROI function to limit the scanning the middle of the target cell’s somata region. The scanning pixel dwell time is increased to 10 μs and various scanning conditions were used in each mouse tested. After the first induction session, the head cover was re-installed to seal the craniotomy from ambient light. Seven to ten days after the first session, all the cells were re-imaged to check tdTomato expression and at the end underwent a 30 minutes blue LED exposure (5 mW, 470 nm, Thorlabs, M470L3). And the third imaging session was carried out seven to ten days after the LED exposure.

For calcium imaging experiment, a custom-made two-photon microscope was used with an objective lens Olympus XLUMPLFLN-W 0.95 NA 20×, 8 kHz resonant scanner head (Cambridge Technologies, CRS8K/6215H). Fluorescence light have been captured using H11706-40 and H10770PA-40 photomultiplier tubes equipped with low noise amplifiers (Femto, DHPCA-100), whose analog signal was subsequently digitized (National Instruments, NI-5732) and formed into images using field programmable gate array (National Instruments, FPGA NI-7961R) and ScanImage software (Vidrio Technologies) [Pologruto et al., Biomedical Engineering Online 2:13, 2003]. GCaMP signal was collected at 30 Hz frame repetition rate for 30 minutes in each mouse.

## Data availability

All relevant plasmids will be deposited to Addgene. Data are available from the corresponding author upon request.

## ACKNOWLEDGMENTS

We are grateful to the Structured Science teams at the Allen Institute for their technical support in stereotaxic injections and mouse colony management. The work was funded by the Allen Institute for Brain Science, NIMH BRAIN Initiative grant RF1MH114106 to A.C., NSFC Science Fund for Creative Research Group of China (Grant No. 61421064) to H.G., Q.L. and S.Z., NIH Director's New Innovator award IDP20D017782 and NIH/NIA 1R01AG047664 to V.G., and Colvin divisional fellowship of Division of Biology and Biological Engineering, California Institute of Technology, to A.K. NIH Brain Initiative U01NS107610 grant to Mark Schnitzer. Immunohistochemistry experiments in Figure 8 were performed in the Biological Imaging Facility, with the support of the California Institute of Technology Beckman Institute and the Arnold and Mabel Beckman Foundation. The creation of Ai139 mouse line was supported by the NIH grant R01DA036909 to B.T. The authors thank Sevi Durdu, Bilal Kerman and Keisuke Yonehara for critical reading and feedback. The authors wish to thank the Allen Institute founder, Paul G. Allen, for his vision, encouragement, and support.

## AUTHOR CONTRIBUTIONS

A.C. conceptualized the light-inducible recombinase system. S.Y. performed cloning and characterization of the constructs as well as participated in image acquisition. B.O. performed all the surgeries and image acquisition.

T.Z. Performed cloning. M.M. performed some of the surgeries and light stimulations. T.D. performed some of the initial cloning experiments. B.T. and H.Z. contributed to the generation of the Ai139 transgenic mice. H.G., Q.L. and S.Z. acquired fMOST data. X.K. and Y.W. performed Neurolucida reconstructions. V.G. and A.K. performed deep brain imaging experiments. S.C. and P.B. performed 2P induced recombination experiments. A.CT. and A.D. performed zebrafish experiments. R.C., P.Y and M.S. performed the targeted single cell 2P experiments and combinatorial cortical GCamp7F calcium imaging experiments. A.C. and H.Z. designed and coordinated the study as well as wrote the manuscript, with inputs from all coauthors.

---

**Supplementary Figure 1.**
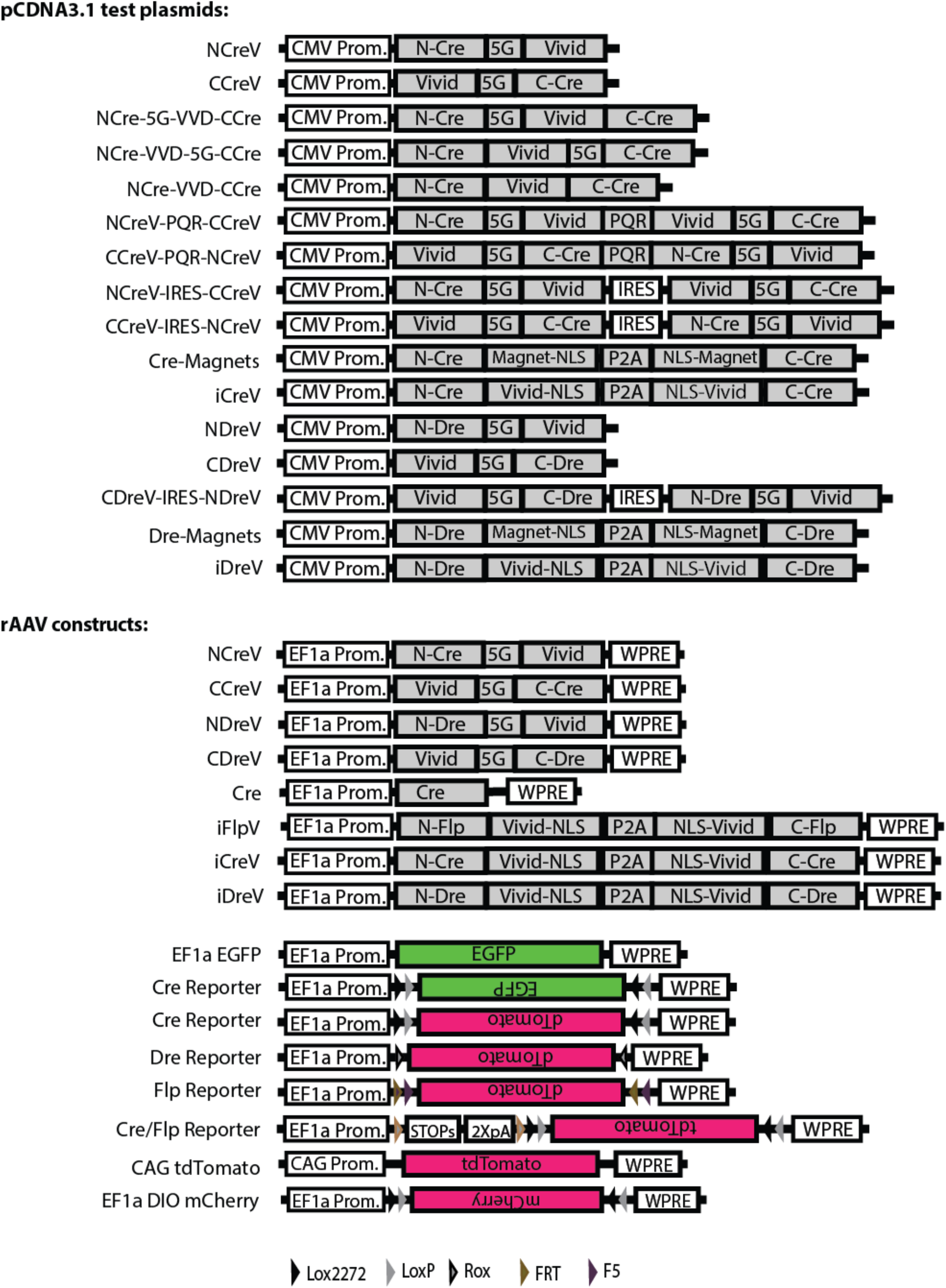
Schematic diagram of DNA constructs generated for the study.

**Supplementary Figure 2.**
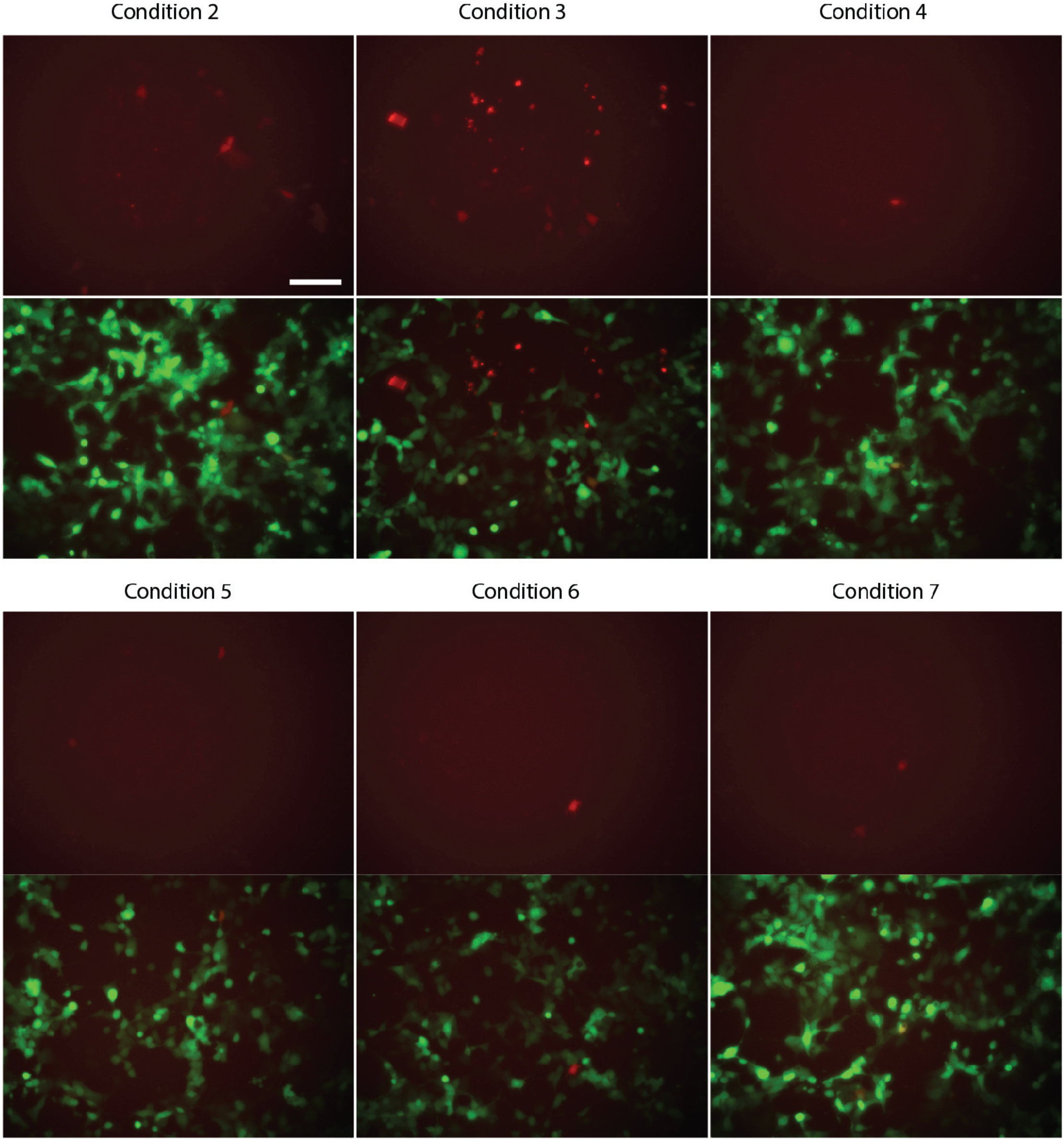
*In vitro* two-photon stimulation of DreV. Mammalian cells were co- transfected with EF1a-EGFP, EF1a Dre red fluorescence reporter, EF1a-NDreV, and EF1a-CDreV plasmids. Two-photon activation was conducted at 48 hr after transfection at various conditions, and reporter expression was observed 36 hr post stimulation. Two-photon activation conditions were as follows: λ = 900 nm, 90 mW, 1 ms/line (512 lines), 200 μm × 200 μm scan area, and duration of: 1) 3 mins (data not shown due to no signal), 2) 6 mins, 3) 9 mins, 4) 12 mins, 5) 12 mins (repeat), 6) 15 mins. In condition 7 randomly selected single cells were scanned in 5 areas, 36 × 36 μm each, separated by 40 μm roughly in a straight line across the plate with a duration of 1-10 seconds per area. Top left scale bar: 100 μm.

**Supplementary Figure 3.**
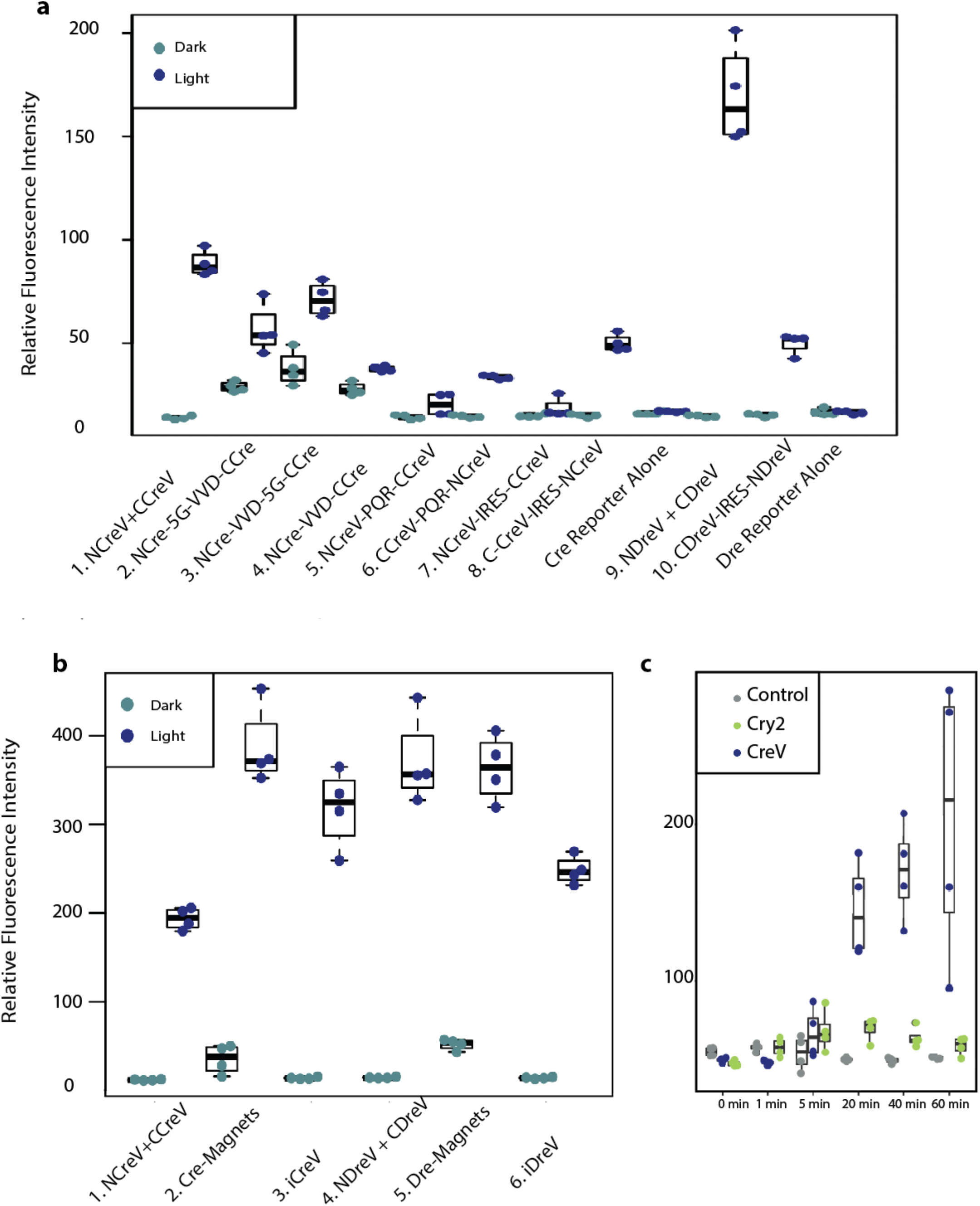
*In vitro* efficiency comparisons of optogenomic modification constructs used in this study. **(a)** Relative fluorescence intensity of Cre dependent and Dre dependent fluorescent reporters 48 hours after 20 minutes of light induction for different co-expression constructs. The N- and the C-termini of the CreV recombinase were combined using a variety of approaches. In the first one, the NCreV and CCreV constructs were mixed together. Constructs 2,3 and 4 contain NCre linked with VVD and CCre all within the same open reading frame, with or without a 5G linker. Constructs 5 and 6 contain both NCreV and CCreV linked by the ribosome skipping peptide PQR. Constructs 7 and 8 have NCreV and CCreV linked by IRES sequence instead of PQR. Constructs 10 is the DreV version of the most successful CreV co-expression construct. Controls are represented by reporters alone. **(b)** Comparison of the light-inducible recombination mediated by improved CreV and DreV. Cells were co-transfected with the appropriate Cre or Dre-dependent reporters, and recombinase constructs 1. NCreV and CCreV, 2. Cre-Magnets 3. iCreV, 4. NDreV and CDreV, 5. Dre-Magnets 6. iDreV. Images taken 48 hours after 20 minutes of light induction. Relative fluorescence intensities were calculated from 4 replicates. **(c)** Comparison of iCreV system with improved CRY2 based light inducible Cre recombination system. Cells were transiently transfected with fluorescent reporter along with either iCreV or improved CRY2/CIB1 based constructs. 48 hours after various durations of light stimulation average fluorescence values were quantified with 4 replicates per condition.

**Supplementary Figure 4.**
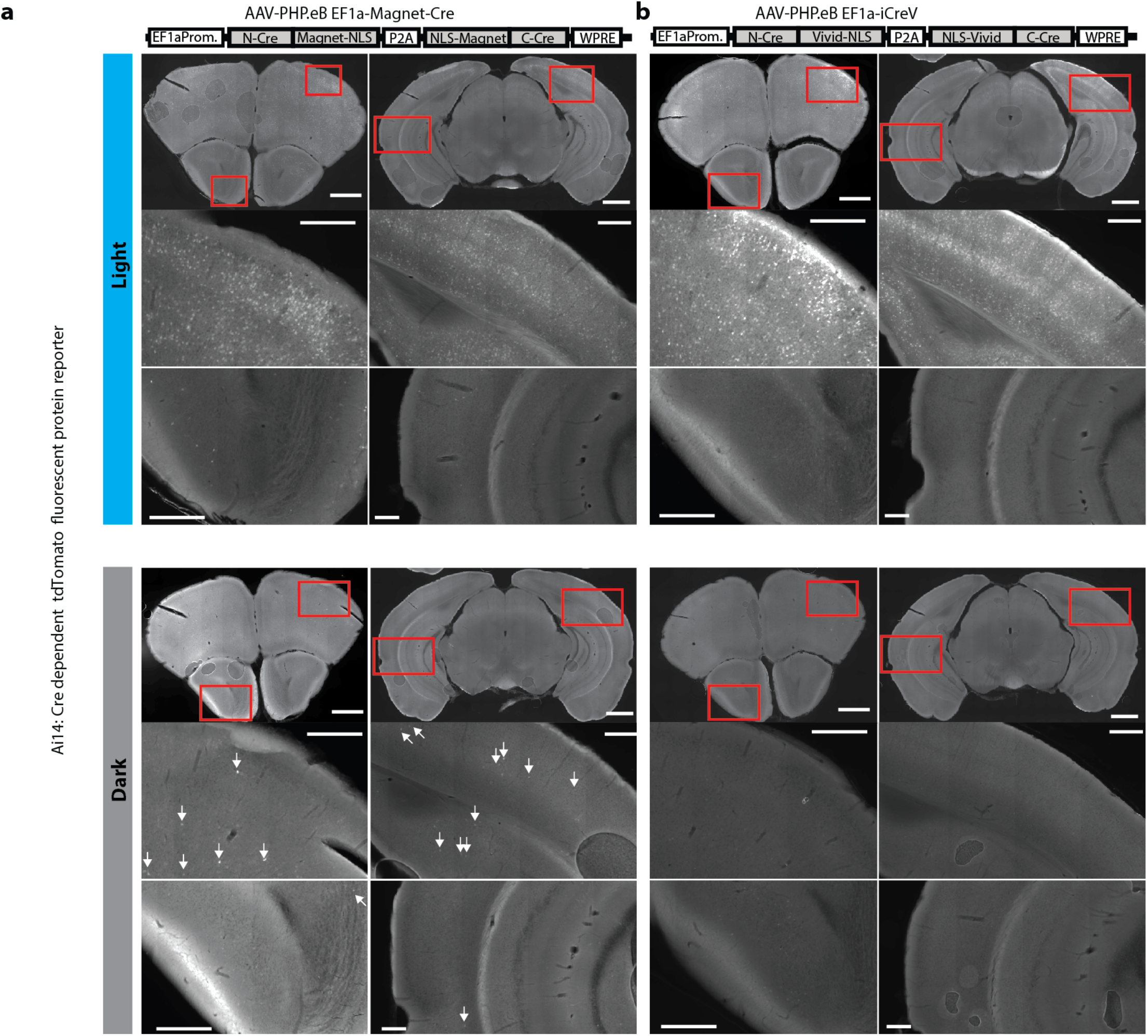
*In vivo* testing and comparison of iCreV with Cre-Magnets. (**a**) Blood brain barrier permeable PHP.eB- iCreV and Cre-Magnet viruses were RO injected along with EF1a-eGFP control viruses into Ai14 mice (n=2 per case). 2 weeks after stimulation 2 mice received light stimulation on the right hemisphere and 2 left as control. Both constructs resulted in robust light induced recombination. In no light cases magnets resulted in consistent recombination throughout the brain indicated by white arrows in contrast with iCreV no light cases. Scale bars: Overviews 1 mm, smaller brain segments 200 μm.

**Supplementary Figure 5.**
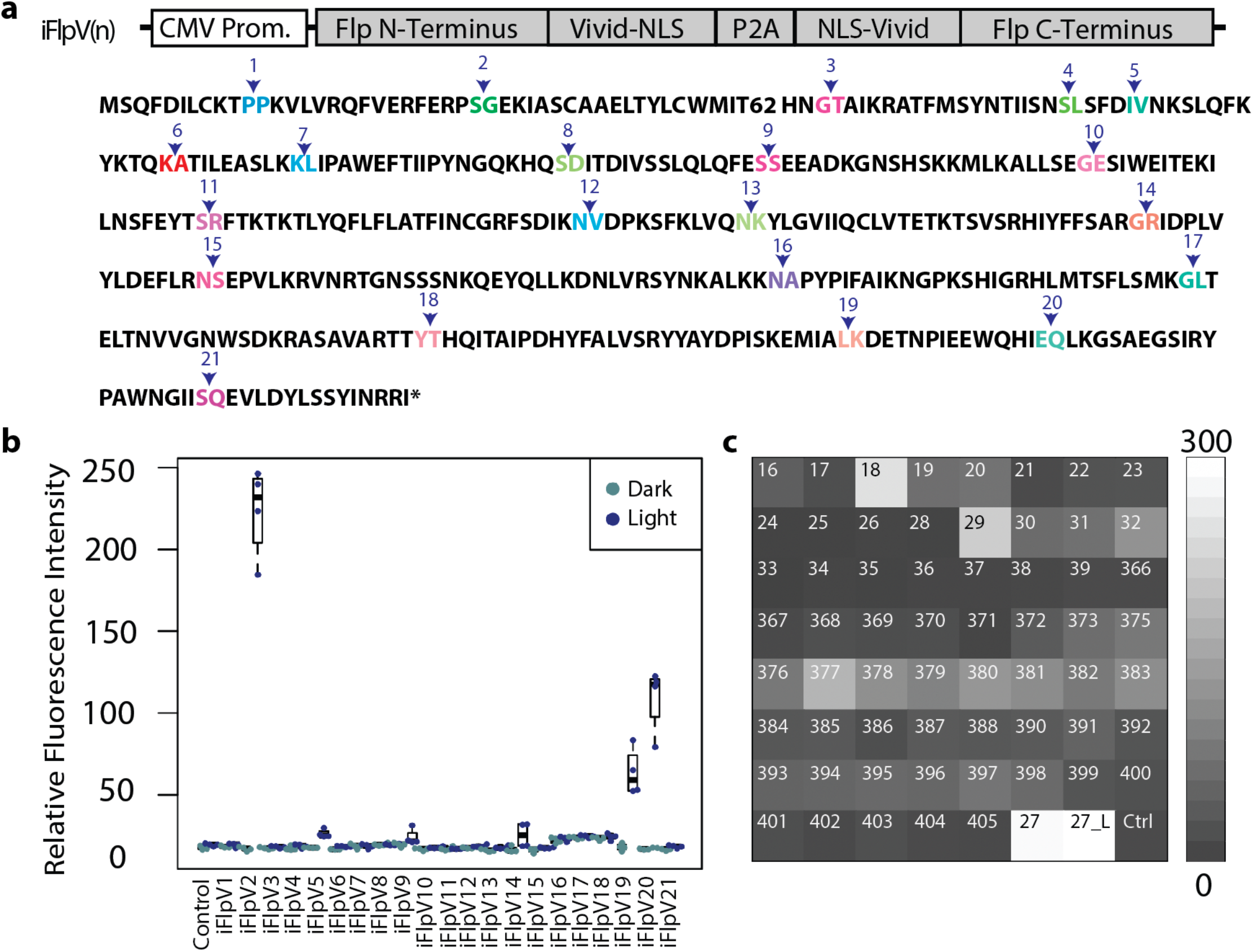
Generation of a light inducible Flp recombinase. (**a**) Initially 21 different iCreV based iFlpV constructs were generated by varying the splitting locations of the Flp open reading frame. (**b**) The expression plasmids containing these constructs were co-transfected into mammalian cells with a Flp-dependent fluorescent reporter, and stimulated with either 20 minutes of light or left in dark (n = 4 per case). 48 hours after light and dark treatments cells were imaged and relative fluorescence intensities were graphed. (**c**) Further screening of 62 iFlpV variants. Split locations are right after the amino acid numbers indicated on the matrix. Mammalian cells were transfected with plasmids expressing iFlpV and its variants and a Flp-dependent fluorescent reporter. Cells were stimulated with either 20 minutes of LED light or left in dark (n=3 per condition). Fluorescence intensity was measured as integrated density in Image J, and the values for the 62 iFlpV variants, iFlpV and negative control were arranged in an 8×8 matrix. The last two brightest conditions are iFlpV2 (aa 27) with an additional linker sequence and iFlpV2 as a control. Overall iFlpV2 resulted in the best induction rate with least background.

**Supplementary Figure 6.**
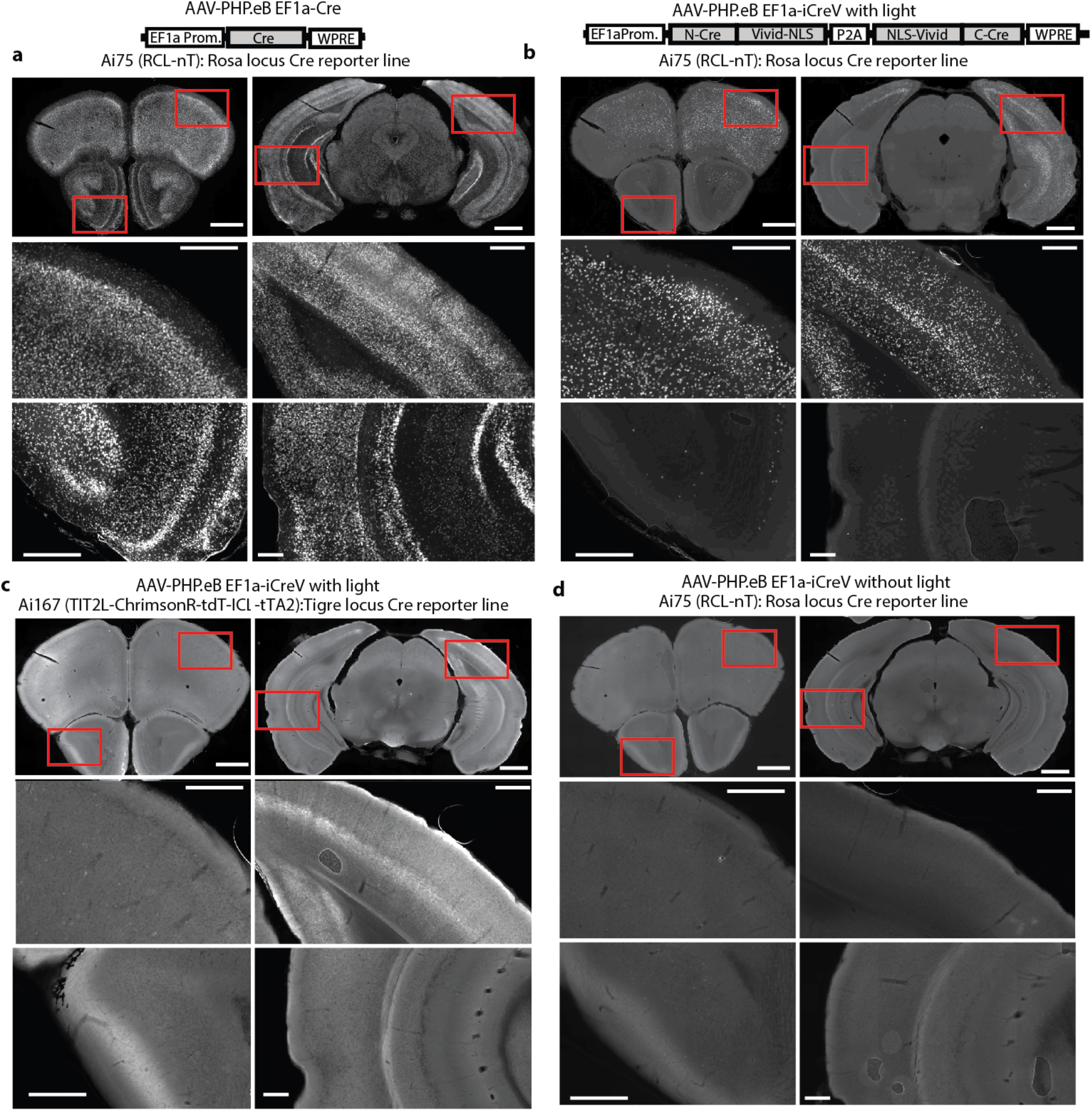
RecV viruses allow efficient light mediated optogenomic modifications at different loci. Various reporter mice (n = 2 per case) each received retroorbital (RO) injection of PHP.eB rAAVs. (**a**) Cre-dependent fluorescent reporter Ai75 mice (from the Rosa26 locus) were injected with AAV-PHP.eB EF1a-Cre virus, leading to widespread recombination. (**b**) Cre-dependent tdTomato-expressing Ai75 mice were RO injected with AAV-PHP.eB EF1a-iCreV viruses. (**c**) Cre-dependent ChrimsonR expressing Ai167 mice (from the TIGRE locus^50^) were RO injected with AAV-PHP.eB EF1a-iCreV. (**d**) Cre-dependent tdTomato-expressing Ai75 mice were RO injected with AAV-PHP.eB EF1a-iCreV viruses and no light was applied. All *in vivo* light activation was applied through the skull on the left hemisphere. Experimental mice were treated with light 2 weeks post virus injection and sacrificed 2 weeks post light stimulation. Two coronal planes are shown for each injection (top row) with enlarged views (lower two rows) for areas indicated by the red boxes. In all experimental cases (**b and c**) light induction led to localized recombination radiating from the light induced hemisphere. When no light was applied in iCreV injected reporter mice there was no significant recombination (c). Scale bars: Overviews 1 mm, smaller brain segments 200 μm.

**Supplementary Figure 7.**
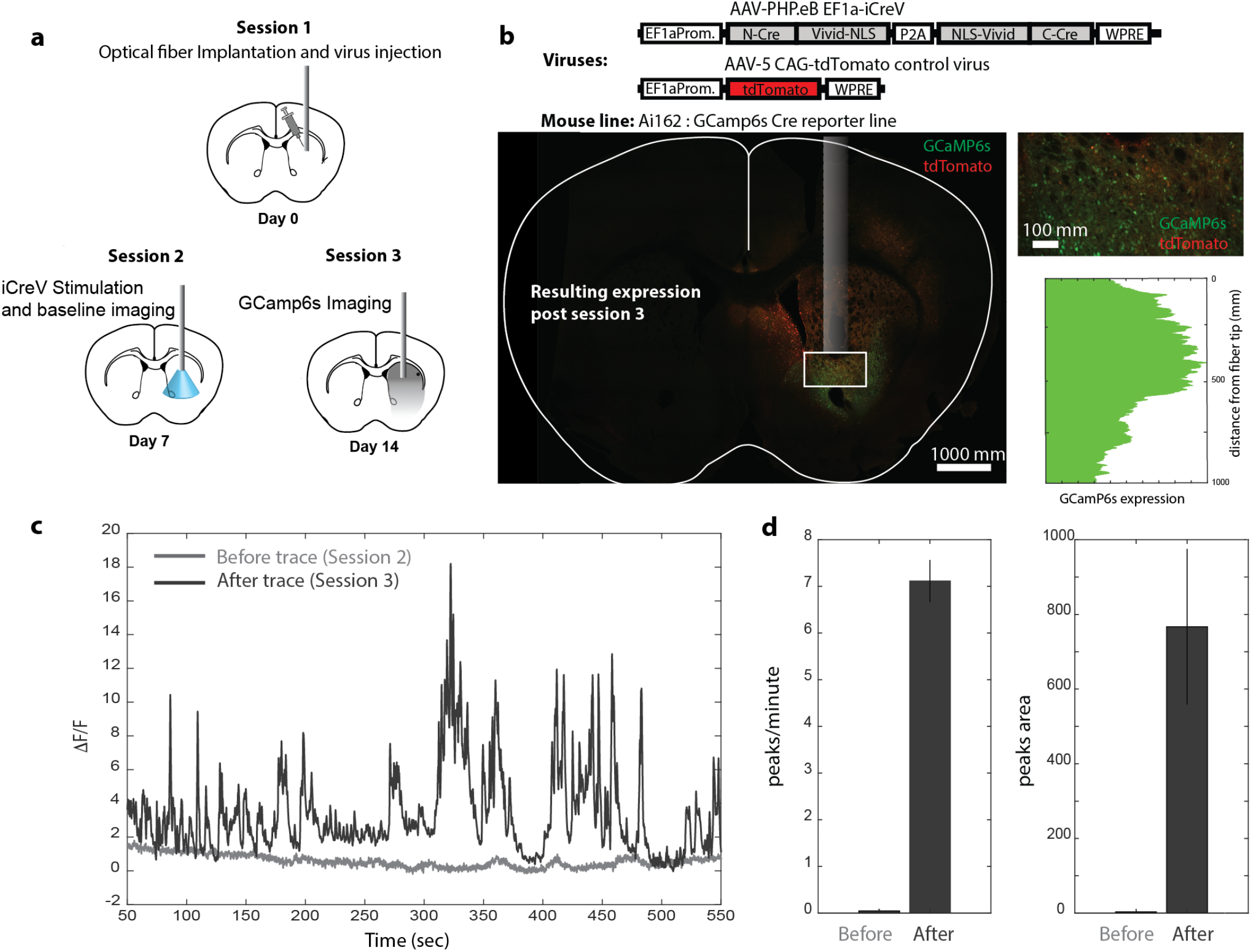
Localized iCreV mediated GCamP6s expression and optical physiology within deep brain structures. (**a**) Virus application scheme and implant for one-photon illumination and GCaMP6s recordings. GCaMP6s reporter mice were locally injected with 1:1 mixture of PHP.eB.iCreV and AAV5.CAG.tdTomato and implanted with 400 μm optical fiber. After 7 days, fiber photometry signal was recorded as baseline activity, and 1P illumination was performed by a 447nm laser using a 200 μm fiber, 5mW, 100ms pulses, 1Hz for 30 minutes. At day 14 fiber photometry signal was measured again. (**b**) Immunohistochemistry showing broad expression of tdTomato, and local and defined expression of GCaMP6s as observed under the fiber tip after iCreV activation. (**c**) Fiber photometry activity in the striatum. ‘Before’ represents baseline activity before illumination -session 2, ‘After’ represents GCaMP6s activity a week after illumination -session 3. (**d**) Plots of ΔF/F over time and area, before and after illumination (n= 2 mice, 1 trial each).

**Supplementary Figure 8.**
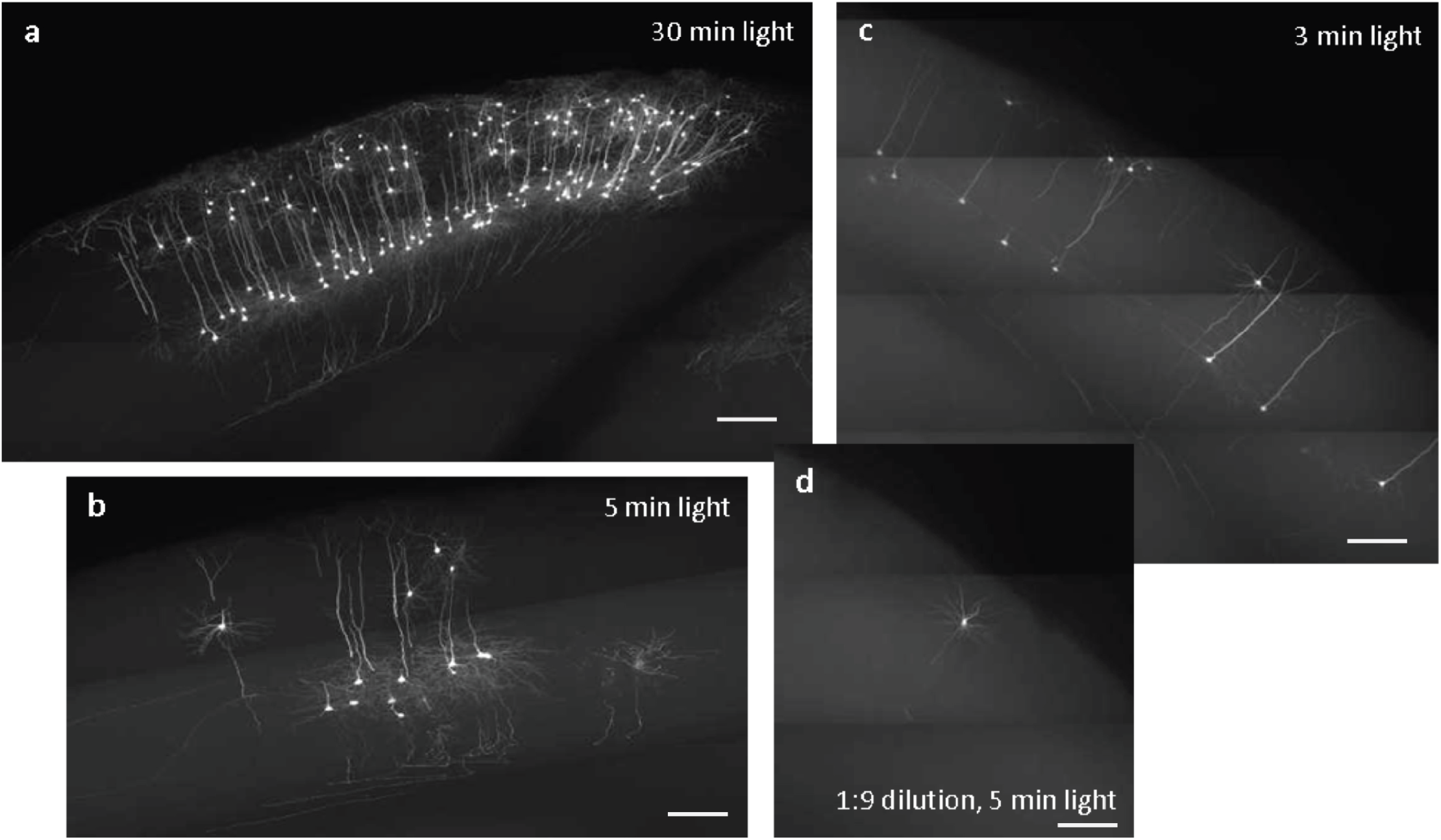
Lower dose of CreV viruses and shorter duration of light induction leads to sparse and strong labeling of individual neurons. Ai139 Cre-dependent EGFP-expressing transgenic mice were injected with the 1:1 mixture of NCreV and CCreV rAAVs in different dilutions in the visual or somatosensory cortex. Two weeks after injection, various durations of light were applied across the skull at the injection sites. Two weeks post light induction, the brains were collected for fMOST imaging. Shown here are examples for the number of neurons labeled at the injection sites: (**a**) injection using undiluted viral solution, 30 minutes of light exposure; (**b**) injection using undiluted viral solution, 5 minutes of light; (**c**) injection using undiluted viral solution, 3 minutes of light; (**d**) injection using 1:9 dilution of viral solution, 5 minutes of light. Images are maximum projections of 100 consecutive fMOST images (each 1 μm-thick), thus equivalent to 100 μm-thick sections. Scale bars: 200 μm.

**Supplementary Figure 9.**
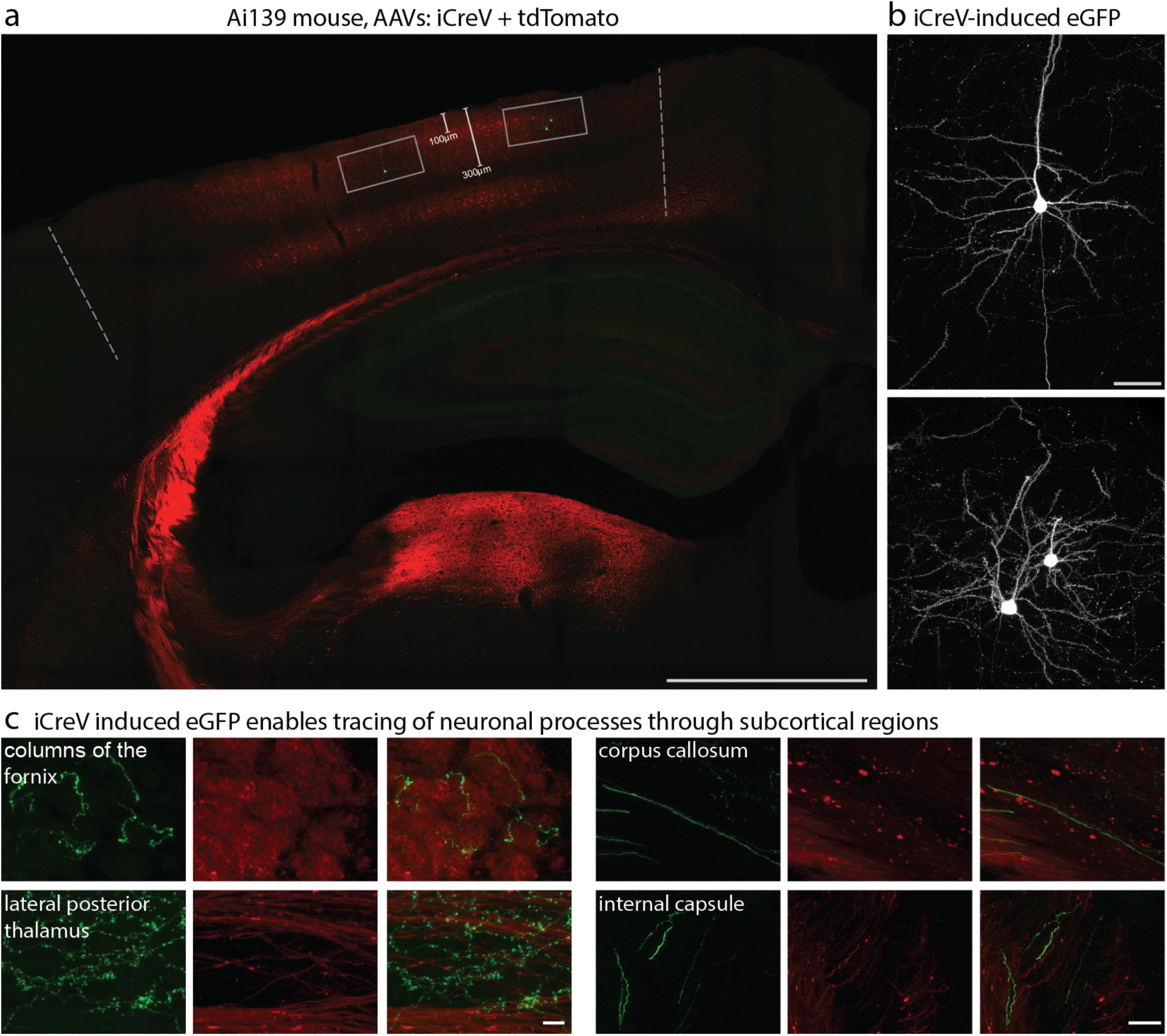
*In vivo* two-photon stimulation and sparse recombination using iCreV. **(a)** Ai139 mice received single stereotaxic injections of a 1:5 mixture of EF1a-iCreV:EF1a-tdTomato into VISp. A glass cranial window and headplate were fitted over the injected region and temporarily covered with Kwik-cast mixed with black powder paint until two-photon stimulation. Gray dotted lines mark edges of the cranial window region. Two to three weeks post-injection, mice were head-fixed while walking on a stable platform, kwik-cast over the cranial window was removed, and discrete 400μm × 400μm regions of layer 2/3, approximately 150-250μm below the pial surface (gray boxes), were stimulated at λ = 910nm for 15 minutes each. Following stimulation, the window was recovered with kwik-cast mixed with black paint and animals were returned to their home cage. Two weeks later, animals were perfused. Scale bar is 1mm. **(b)** High magnification images of eGFP-filled neurons within stimulated regions in (a). Two-photon stimulation of iCreV induced sparse eGFP expression within each region, visualizing individual neurons and their associated processes. Scale bar is 50μm. (**c**) Individual neuronal processes (right panels) and terminations (left panels) are traceable through multiple subcortical regions, enabling brain-wide morphological reconstructions of individual neurons. Scale bars are 10μm (left) and 30μm (right).

**Supplementary Figure 10.**
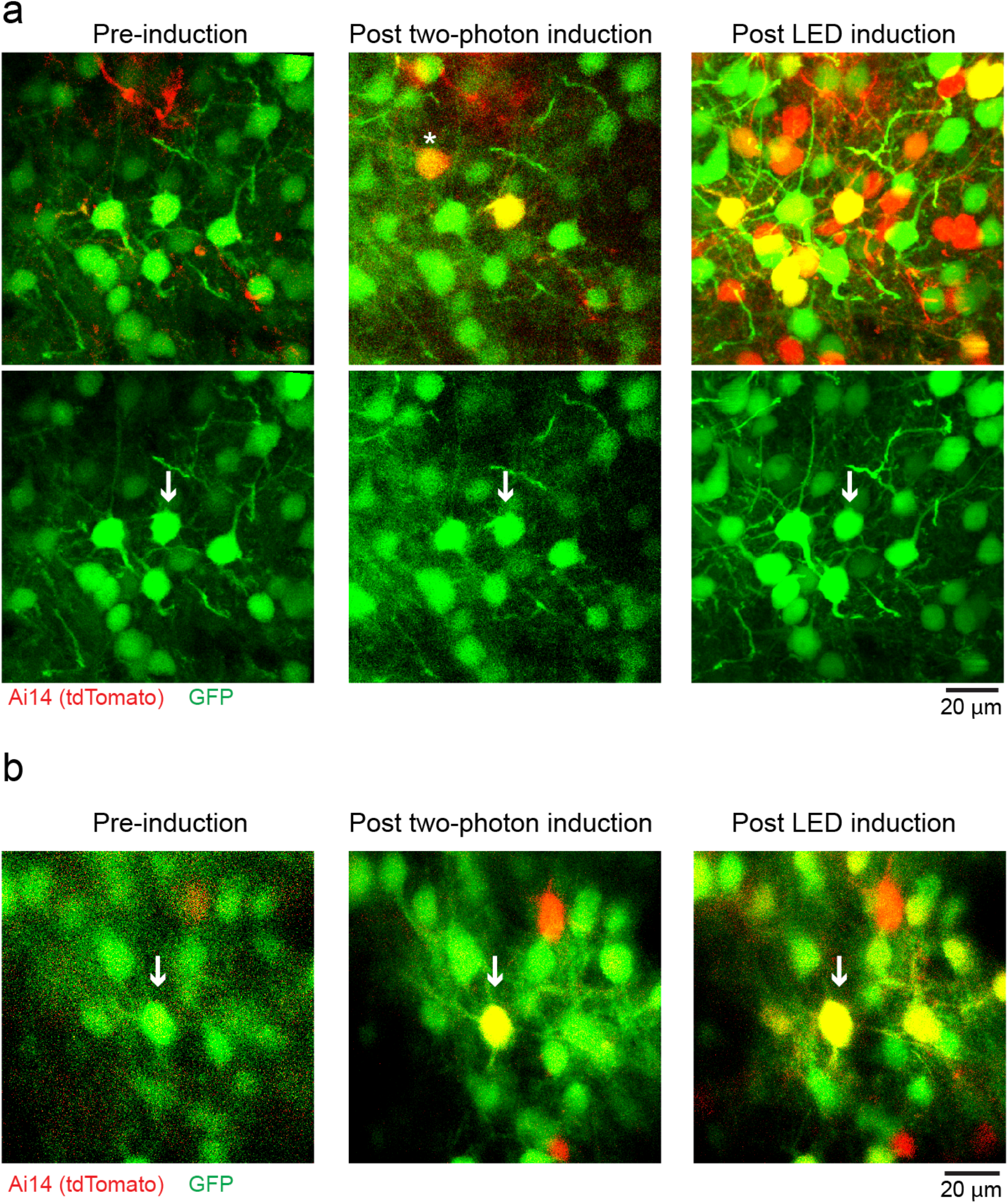
Additional examples of the in vivo single cell 2P induction experiment. Additional two examples (a, b) of the 2P induction experiment, following the same preparation and experiment protocol in **Figure 5b**. Arrows indicate the target cells. Asterisk indicates an induced non-targeted cell. The background red signal observed in the pre-induction session is due to either incomplete shielding of ambient light after surgery or high multiplicity of infection related issues. See Supplementary Figure 11 for background expression rate.

**Supplementary Figure 11.**
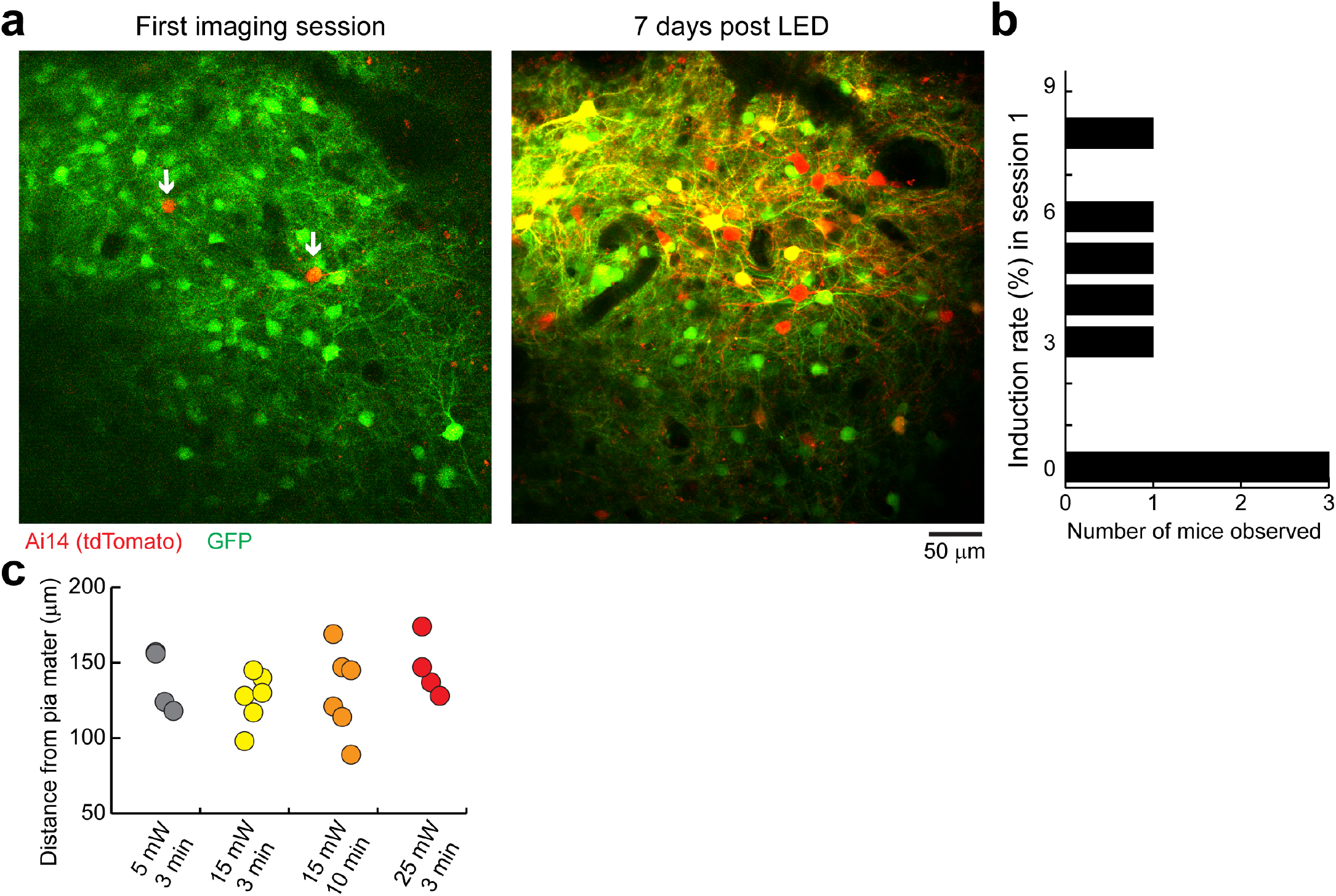
Background induction rate observed in the 2P experiment mice cohort. (a)Example images of the first imaging session showed some cells already expressed tdTomato, indicating some background induction prior to two-photon stimulation. Arrows indicate the induced cells. (**b**) Quantification of this background induction rate showed an average of 3% (N = 9 mice). This induction was likely due to incomplete shielding of the ambient light or high multiplicity of infection. (**c**) Quantification of all the induction plane depths for each field of view associated with the two-photon induction experiment (related to **Figure 5c** and **d**).

